# Computational Resilience in Human Reciprocity: An Asymmetric Intrinsic Prosocial Bias for Sustaining Cooperation under Exogenous Uncertainty

**DOI:** 10.1101/2025.05.03.651994

**Authors:** Xuqi Liu, Chi Yao, Rui Liao, Yu Nan, Yuankun Fang, Yang Hu, Xiaolin Zhou, Xiaoxue Gao

## Abstract

In direct reciprocity, environmentally imposed exogenous uncertainty frequently decouples benefactor’s altruistic intentions from final outcomes. How do cooperative bonds remain resilient against this inevitable “noise”? Adopting a comprehensive levels-of-analysis framework across eight experiments, we integrate interpersonal paradigms, computational modeling, multivariate fMRI decoding, and third-party social evaluations to uncover a robust “asymmetric adjustment” in beneficiary’s affective evaluation and reciprocity under exogenous uncertainty, systematically delineating its cognitive algorithmic rules, neural representations, and adaptive functions in sustaining human cooperation. Specifically, beneficiary’s reciprocity amplifies for positive deviations from expectations while buffering against reciprocal reductions from unintended shortfalls. Modeling reveals this computational resilience is governed by an asymmetric intrinsic prosocial bias, extending beyond traditional frameworks of outcome/intention-based evaluation, prediction-error-based social learning, or loss aversion. Supported by theory-of-mind system, this “asymmetric adjustment” is perceived as morally superior to other calculations, thereby serving a critical adaptive function to sustain cooperation amidst the volatility of an uncertain world.

## Introduction

Direct reciprocity, whereby beneficiaries evaluate and reciprocate benefactors’ altruistic acts, is fundamental to human cooperation and adaptation^1–6^. Real-world direct reciprocal interactions are often embedded in dynamic and fluctuating uncertainty. Previous studies have predominantly focused on endogenous uncertainty, which resides in a beneficiary’s beliefs about the benefactor’s intentions and actions (e.g., whether and to what extent the benefactor will help)^7,8^. Beneficiaries can proactively resolve such uncertainty through interpersonal interactions and social learning, thereby updating their expectations with the benefactor’s actual intentions and behaviors ^9–12^ and adjusting their affective evaluations (e.g., gratitude) and reciprocal behaviors accordingly^13–15^. In an “ideal social world”, this synergistic interplay between belief updating and the gradual resolution of endogenous uncertainty can ultimately foster a stable and predictable social bond between the two parties^7,16,17^.

However, real-world cooperative relationships are rarely noise-free or permanently static once expectations are formed. They are frequently challenged by exogenous uncertainty, i.e., external volatility that is often beyond the control of both parties and can inevitably decouple an actor’s intentions from the final outcomes^8,18,19^. For example, during a natural disaster, the actual cost a benefactor incurs to rescue a beneficiary may ultimately turn out to be higher or lower than both parties initially expected. How do a beneficiary’s affective evaluations (e.g., gratitude) and reciprocity dynamically adjust when transitioning from exogenous uncertain expectations to better or worse actual outcomes? How do human social bonds built on direct reciprocity remain resilient in face of the inevitable “noise” arising from exogenous uncertainty? Understanding these questions is crucial for comprehending and predicting human reciprocity, cooperation and adaptation, which is essential for guiding daily social decision-making^7,12,20^ and informing policy aimed at fostering cooperation^21–25^. Nevertheless, given that prior research has extensively focused on either the non-dynamic events in certain contexts^20,26–35^ or the dynamic reduction of endogenous uncertainty through social learning^7,9–15^, the cognitive and neural mechanisms underlying how humans navigate and buffer against exogenous uncertainty in direct reciprocity remains largely unexplored.

While existing studies of gratitude and reciprocity do not explicitly capture these exogenous dynamics, they lay the conceptual groundwork for investigating this adaptation. Within certain and static environments, extensive research establishes that individuals evaluate altruism through intention-based and outcome-based mechanisms^3,28,29,34–40^. These dimensions also serve as vital cognitive antecedents when transitioning from exogenous uncertainty to certainty. During the uncertain phase, beneficiaries form expectations regarding the benefactor’s intended cost, reflecting their benevolent intention. Conversely, the actual cost ultimately borne, which remains beyond either party’s control, constitutes the final outcome. If beneficiaries rely predominantly on intention-based evaluation^3,28,34,37,38,41^, their gratitude and reciprocity should remain anchored to the initial expectations and unchanged by the actual outcome. Conversely, an outcome-based evaluation^3,28,35,37,42^ predicts that reciprocity will vary in accordance with the final realized cost. Complementing these static accounts, dynamic models of social interaction suggest that beneficiaries responds to endogenous uncertainty fluctuations in beliefs about the benefactor’s intentions and actions via symmetrical prediction-error-based social learning^13^. Under this framework, beneficiaries should integrate prior expectations with final outcomes and update their responses based on the discrepancy between the two (i.e., prediction error, PE) linearly and symmetrically for positive and negative PEs. Consequently, gratitude and reciprocity should adjust linearly and symmetrically, scaling precisely with the magnitude of the discrepancy between the expected and actual costs.

Crucially, aforementioned prevailing theories on gratitude and reciprocity, whether focusing on the outcome-based evaluation, intention-based evaluation, or symmetrical PE-based social learning, consistently predict a symmetric change in gratitude and reciprocity when the final certain outcome turns out to be better or worse than what was expected under exogenous uncertainty. However, employing a multi-round interpersonal task simulating the transition from exogenous uncertainty to certainty (Fig. 1; adapted from Xiong et al., 2020^41^), we uncover a novel “asymmetric adjustment” that challenges these accounts. Specially, in each round of the task (taking Experiment 1 as example; see *Methods*), participants were paired with an anonymous co-player, who decided whether to endure an exogenous uncertain cost (with a 50% chance of being either 2 or 8 times, determines randomly by the computer system, expected to be 5 times of pain stimulation) to reduce the participant’s pain by half (from 10 to 5 times). After presenting the co-player’s decision, participants learned the co-player’s actual incurred cost, which was either higher (i.e., 8 times) or lower (2 times) than the expected. We quantified participants’ gratitude via trial-by-trial subjective ratings during the task both before and after the resolution of this uncertainty. Results demonstrated that, the beneficiary’s gratitude intensified significantly when the final benefactor-cost exceeds the expectation under exogenous uncertainty; however, a parallel reduction in benefactor-cost did not elicit equivalent decrease.

**Fig. 1.**
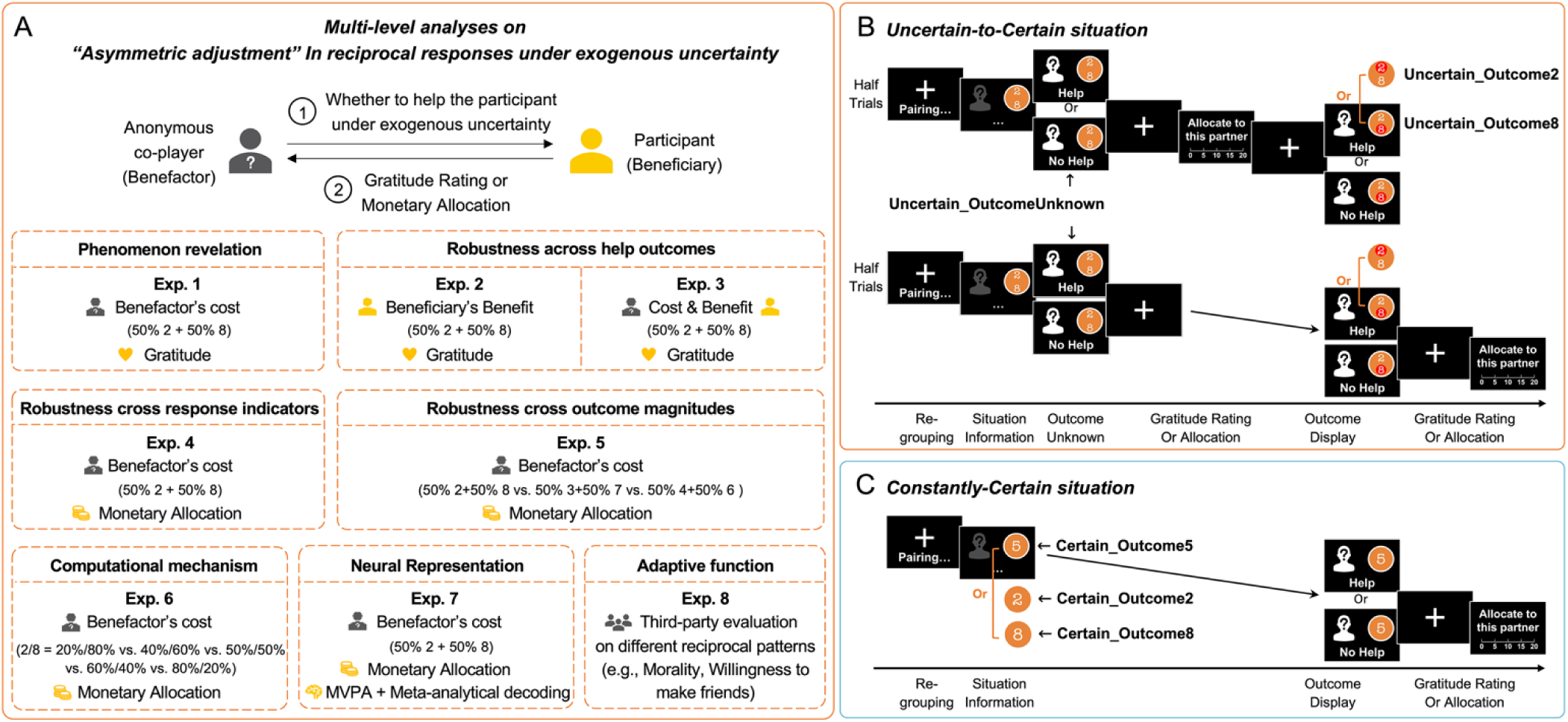
Multi-level analyses framework and experimental procedures. **(A)** Overview of the study’s multi-level framework across eight experiments, investigating the “asymmetric adjustment” in direct reciprocity under exogenous uncertainty. These experiments were designed to delineate the phenomenon at complementary dimensions: its fundamental behavioral manifestation (Experiments 1–5), its governing cognitive computational rules (Experiment 6), its biological implementation in the brain (Experiment 7), and its adaptive social function (Experiment 8). **(B)** Procedure for the Uncertain-to-Certain situation. In each round, the participant (beneficiary) was paired with an anonymous co-player (benefactor). The co-player decided whether to help reduce the participant’s noise/pain stimulation under an exogenous uncertain cost (e.g., 50% probability of 2 or 8 times in Experiments 1-4 and 7). Participants reported their gratitude to the co-player (Experiments 1-3) or made a monetary allocation between self and the co-player (Experiments 4-7) either before the actual cost was revealed (Uncertain_OutcomeUnknown) or after observing a higher or lower actual cost (e.g., Uncertain_Outcome8 or Uncertain_Outcome2). **(C)** Procedure for the Constantly-Certain situation. The benefactor’s cost was certain from the outset (e.g., 2, 5, or 8 in Experiments 1-4 and 6-7), serving as a baseline to isolate the cognitive processing specific to the uncertain-to-certain transition.

To elucidate the cognitive mechanisms driving this “asymmetric adjustment”, we draw upon broader theoretical frameworks and empirical evidence from behavioral economics, social cognition, and evolutionary psychology that extend beyond the specific contexts of gratitude and reciprocity. By integrating these perspectives, we propose three competing cognitive hypotheses:

1. *The “Loss Aversion” Hypothesis*. Drawing on prospect theory from economics, individuals are generally more sensitive to losses than to equivalent gains^43–45^. Thus, a higher-than-expected cost incurred by the benefactor may be perceived as a greater “loss”, exerting a disproportionately stronger influence on the beneficiary’s reciprocal responses compared to the equivalent “gain” of a lower-than-expected cost.
2. *The “Differential PE Sensitivity” Hypothesis*. Research across social and non-social learning domains suggests that individuals often update their beliefs differently in response to positive versus negative prediction errors^46–48^. Consequently, the “asymmetric adjustment” may stem from a disparity in how beneficiaries dynamically update their beliefs when outcomes are better or worse than expected.
3. *The “Intrinsic Prosocial Bias” Hypothesis*. Grounded in error management theory^49^, cognitive biases in perception, attention, and social judgments under uncertainty may have evolved as adaptive responses to the asymmetries in the costs of false-positive and false-negative errors throughout evolutionary history; selection should have favored a bias toward making the least costly error. In the context of sustaining cooperation, the asymmetries in the costs of false-positive and false-negative errors also exist. For example, the cost of mistakenly dissolving a relationship with a high-quality cooperator by reducing reciprocity due to environmental noise (a false negative) commonly outweighs the cost of occasionally over-reciprocating (a false positive). Therefore, it is possible that the “asymmetric adjustment” in reciprocity observed under exogenous uncertainty may be driven by an intrinsic prosocial bias that serves to minimize overall social costs and sustain human cooperation.

To distinguish between these competing hypotheses and advance a holistic, mechanistic understanding of this “asymmetric adjustment,” we structured our investigation upon a levels-of-analysis framework^50,51^. This framework emphasizes the critical importance of systematically integrating the investigations on the adaptive social function (Marr’s computational level), the underlying cognitive computations and rules (Marr’s algorithmic level), and the biological implementation in the brain (Marr’s implementational level) to understand social behaviors or emotions. Because these three levels are highly interdependent and mutually constraining, investigating any single level in isolation risks yielding a fragmented explanation that misses the broader theoretical forest. Specifically, to establish the empirical foundation for this multi-level analyses (Fig. 1A), we first conducted a series of five experiments examining the fundamental manifestation and robustness of the “asymmetric adjustment.” After uncovering the phenomenon (Experiment 1), we examined its robustness across distinct help-outcome structures (Experiments 2 & 3: benefactor-cost vs. beneficiary-benefit), established its extension to objective and incentivized behavioral metrics (Experiment 4: monetary allocations), and parametrically evaluated its stability across varied outcome magnitudes (Experiment 5).

Having firmly established this behavioral target, we proceeded to reveal its underlying mechanisms from the aforementioned three levels. First, at the cognitive computation level, we sought to identify the latent rules generating the “asymmetric adjustment.” By systematically modulating prior probabilities to decouple outcome magnitudes from prediction errors, we applied formal computational modeling to fit and compare competing mathematical models (Experiment 6), dissociating whether this “asymmetric adjustment” is governed by differential sensitivity to positive vs. negative PEs or the intrinsic prosocial bias. Second, at the biological implementation level, to provide neural evidence supporting these computationally identified cognitive mechanisms, we integrated the interpersonal task with functional magnetic resonance imaging (fMRI) multivariate pattern analysis (MVPA) and meta-analytical neural decoding (Experiment 7) to pinpoint the neural representations supporting these dynamic computations. Finally, drawing upon the mechanistic clues provided by the cognitive and biological levels, we formulated and tested a hypothesis regarding the adaptive function of this behavior, which reciprocally enriches our understanding of why the human brain favors these specific cognitive and neural foundations. Finally, building upon the mechanistic clues provided by the cognitive computation and biological implementation levels, we hypothesized that the “asymmetric adjustment” serves a crucial adaptive function in promoting cooperation, and empirically tested this hypothesis using a third-party social evaluation paradigm (Experiment 8). In turn, understanding this adaptive function reciprocally enriches our comprehension of why these specific cognitive and biological foundations evolved. By spanning these three complementary levels, our article provides a comprehensive account of human direct reciprocity under environmental noise.

## Results

### Uncovering the behavioral manifestation of “asymmetric adjustment” under exogenous uncertainty

We first characterized the fundamental behavioral manifestations of dynamic adjustments in beneficiary’s gratitude and reciprocity under environmental noise. In Experiment 1, we established the core interpersonal paradigm to determine whether and how beneficiaries dynamically adjust their gratitude and reciprocity when a benefactor’s actual cost transitioning from exogenous uncertain expectations to better or worse actual outcomes.

In each round of the interpersonal task of Experiment 1 (Fig. 1B; see details in *Procedures*), participants were paired with a different anonymous co-player, who decided whether to endure an amount of noise (i.e., benefactor’s cost) to reduce the participant’s noise by half (from 10 to 5 times, 1 time lasted 5 seconds). In the Uncertain-to-Certain situation, the co-player made altruistic decision under an exogenous uncertain cost (with a 50% chance of being either 2 or 8 times, determined randomly by the computer system, expected to be 5 times of noise stimulation). After presenting the benefactor’s decision to help under exogenous uncertainty during the Outcome_Unknown phase (Uncertain_OutcomeUnknown condition), we additionally included an Outcome_Display phase, where participants learned the co-player’s actual cost, which was either 8 (higher than the expected 5; Uncertain_Outcome8 condition) or 2 (lower than the expected 5; Uncertain_Outcome2 condition). We additionally included the Constantly-Certain situation as the control, where the benefactor’s cost was constantly certain, with 2, 5 or 8 times (Certain_Outcome2, Certain_Outcome5, or Certain_Outcome8 conditions). We collected participants’ trial-by-trial gratitude ratings for each of the above six conditions.

First, to reveal whether and how beneficiaries dynamically adjusts their gratitude ratings from facing an exogenous uncertain outcome (i.e., benefactor’s cost with a 50% chance of being either 2 or 8, expected to be 5 times) to facing a final actual outcome of higher (i.e., 8) or lower (i.e., 2) benefactor’s cost than expected, we conducted a one-way repeated-measure analysis of variance (ANOVA) and observed a significant main effect of Outcome in the Uncertain-to-Certain situation (Uncertain_Outcome2 vs. Uncertain_OutcomeUnknown vs. Uncertain_Outcome8) (*F*(1.43, 45.69) = 43.83, *p* < 0.001, *η*^2^ =0.58; Fig. 2A; Table S1). Consistent with previous evidence regarding the effect of benefactor’s cost on beneficiary’s gratitude and reciprocity^28,35^, participants’ gratitude ratings (*t*(32) = 7.36, *p* < 0.001, Cohen’s *d =* 1.60) were significantly higher when the actual benefactor’s cost was 8 times than when it was 2 times. Moreover, from the perspective of Uncertain-to-Certain transitions, compared with the Uncertain_OutcomeUnknown condition, participants’ gratitude ratings significantly increased in the Uncertain_Outcome8 condition (*t*(32) = 6.18, *p* < 0.001, Cohen’s *d* = 1.06), and significantly decreased in the Uncertain_Outcome2 condition (*t*(45) = -4.56, *p* < 0.001, Cohen’s *d* = *-*0.54).

**Fig. 2.**
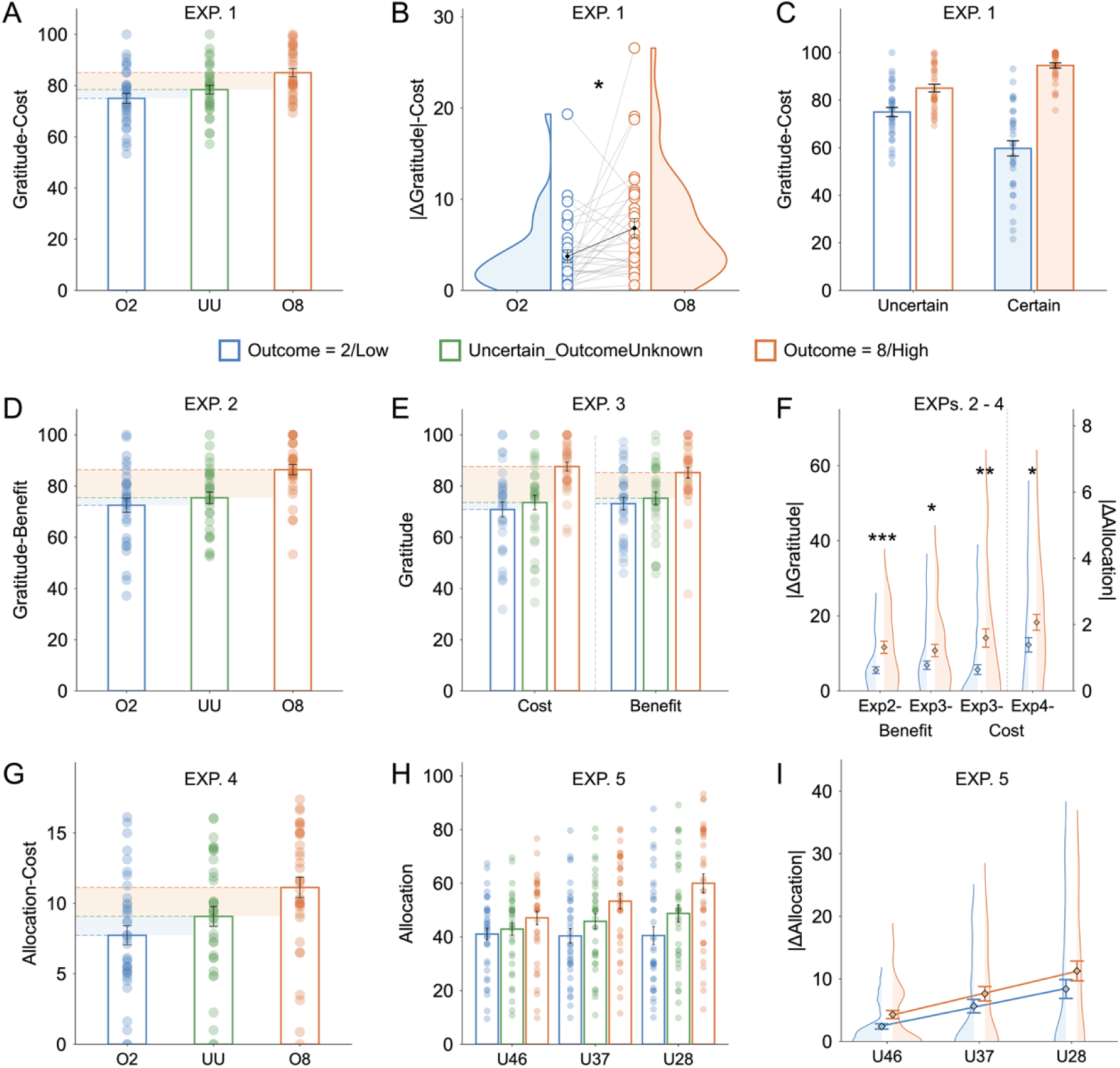
Uncovering and establishing the robustness of “asymmetric adjustment” in reciprocity under exogenous uncertainty. **(A)** Beneficiary’s gratitude ratings to the co-players for different conditions (O2: Outcome = 2; UU: Uncertain_OutcomeUnknown; O8: Outcome = 8) in the Uncertain-to-Certain situation in Experiment 1 (*n* = 33). **(B)** The absolute magnitude of adjustments in gratitude (|△ Gratitude|) when transitioning from uncertainty to an actual outcome of 2 or 8 in Experiment 1. The adjustment was significantly larger when the outcome exceeded expectations than when it fell short. **(C)** Comparison of gratitude ratings between the Uncertain-to-Certain and Constantly-Certain situations in Experiment 1, demonstrating altered outcome processing during the uncertainty resolution. **(D-E)** Robustness of the “asymmetric adjustment” across help outcomes: gratitude ratings when manipulating exogenous uncertainty in the beneficiary’s benefit (Exp. 2, *n* = 31) and both in the benefactor’s cost and in the beneficiary’s benefit (Exp. 3, *n* = 33). **(F)** The “asymmetric adjustment” effect across Experiments 2-4, persisting across help types and response indicators. **(G)** The “asymmetric adjustment” persisted when using monetary allocations reflecting objective reciprocity as the response indicator in Experiment 4 (*n* = 37). **(H)** Robustness of the “asymmetric adjustment” across outcome magnitudes: monetary allocations under 4-or-6, 3-or-7, and 2-or-8 uncertain benefactor-cost conditions in Experiment 5 (*n* = 38). **(I)** Although the outcome magnitude parametrically modulated the absolute degree of change in allocations before and after the resolution, the “asymmetric adjustment” robustly persists across various outcome magnitudes, and its effect size is not significantly modulated by the absolute scale of the outcome deviation. Error bars represent standard errors. \**p* < 0.05, \*\**p* < 0.01, \*\*\**p* < 0.001.

Next, we investigated whether transitions from Uncertain_OutcomeUnknown condition to Uncertain_Outcome8 condition and to Uncertain_Outcome2 condition, both involving an identical absolute change in the benefactor’s cost (i.e., |8 - 5| = |2 - 5| = 3 times), exhibited similar or different magnitudes of influences on the dynamic adjustments in the beneficiary’s gratitude? To answer this question, we subtracted the gratitude ratings in the Uncertain_Outcome2 and Uncertain_Outcome8 conditions from those in Uncertain_OutcomeUnknown condition respectively, and took the absolute values as the indicators of the extent of dynamic adjustments. Paired-samples *t*-test showed that, when the final actual benefactor’s cost was ascertained to be 8 times, which was higher than expected, the extent of adjustments in gratitude ratings were significantly larger than those when benefactor’s cost was ascertained to be 2 times, which was lower than expected (*t*(32) = 2.65, *p* = 0.012, Cohen’s *d =* 0.46; Fig. 2B; Table S2). These findings demonstrated an “asymmetric adjustment”: the beneficiary’s gratitude intensified when the benefactor’s final actual cost exceeded expectations; however, a parallel reduction in the benefactor’s cost did not elicit equivalent decreases.

One plausible explanation is that in the Uncertain-to-Certain situation, the transition from exogenous uncertainty to certainty fundamentally alters the beneficiary’s cognitive processing of the outcome during the final resolution phase, relative to the Constantly-Certain situation where the outcome of help is ascertained from the outset. To test this possibility, we examined whether and how the influence of actual outcomes of the benefactor’s cost (2 vs. 8) on the beneficiary’s gratitude differ across Uncertain-to-Certain situation and Constantly-Certain situation, despite the final outcomes being equivalent in both situations. To this end, we combined the data of the Certain_Outcome2, Certain_Outcome8, Uncertain_Outcome2 and Uncertain_Outcome8 conditions for gratitude ratings, and conducted 2 (Situation: Uncertain-to-Certain vs. Constantly-Certain) × 2 (Actual Outcome: 2 vs. 8) ANOVA (Fig. 2C; Table S3). Results revealed significant main effects of Actual Outcome (*F*(1, 32) = 134.17, *p* < 0.001, *η*^2^ = 0.81) and Situation (*F*(1, 32) = 5.60, *p* = 0.024, *η*^2^ = 0.15). Post-hoc comparisons revealed that gratitude ratings in the Outcome8 conditions was significantly higher than in the Outcome2 conditions (*t*(32) = 11.58, *p* < 0.001, Cohen’s *d* = 1.80), and these in the Uncertain-to-Certain situation was significantly higher than in the Constantly-Certain situation (*t*(32) = 2.37, *p* = 0.024, Cohen’s *d* = 1.80). Importantly, we observed significant interaction effects between Situation and Actual Outcome (*F*(1, 32) = 83.40, *p* < 0.001, *η*^2^ = 0.72). Specifically, participants reported significantly higher levels of gratitude to the co-player when the benefactor’s cost was high (i.e., 8) compared to when it was low (i.e., 2) in the Constantly-Certain situation (*t*(32) = 11.41, *p* < 0.001, Cohen’s *d =* 2.80); these effects were significantly reduced in the Uncertain-to-Certain situation (*t*(32) = 7.36, *p* < 0.001, Cohen’s *d =* 0.81). These results indicated that the contributions of the benefactor’s cost to gratitude were reduced in the Uncertain-to-Certain situation compared to the Constantly-Certain situation, suggesting that there may be different cognitive processes of outcomes involved in the two situations.

### Establishing the robustness of the “asymmetric adjustment” across different types of help outcomes, response indicators and magnitudes of outcome

Having uncovered the basic “asymmetric adjustment” phenomenon in Experiment 1, we next conducted four subsequent experiments (Experiments 2 to 5) to systematically examined whether this phenomenon robustly persists, maintaining a consistent directional asymmetry, across different types of help outcomes (benefactor-cost versus beneficiary-benefit), response indicators (gratitude versus monetary reciprocity), and outcome magnitudes.

From the perspective of the types of help outcomes, we manipulated exogenous uncertainty in beneficiary-benefit instead of or in addition to benefactor-cost in Experiment 2 and 3. The procedures of these two behavioral experiments were similar as Experiment 1, except that in Experiment 2, we manipulated the exogenous uncertainties in the participants’ own benefits instead of that in the benefactor’s cost, while in Experiment 3, we included two blocks, manipulating the exogenous uncertainties in the benefactor’s cost and in the participants’ own benefits, respectively (see *Methods*). The results of Experiment 2 and Experiment 3 revealed the same pattern of asymmetric dynamic adjustments in beneficiary’s gratitude as observed in Experiment 1, not only for the benefactor’s cost, but also for the beneficiary’s benefit, with no significant difference between these two types of manipulations (Fig. 2D-F; Tables S1 and S2). These results not only confirm the robustness of the “asymmetric adjustment” across diverse help-outcome contexts but also demonstrate that this effect cannot be accounted for by the “Loss Aversion” Hypothesis. Specifically, according to the loss aversion mechanism in prospect theory, the results of the beneficiary-benefit manipulations should oppose those of Experiment 1, given that a higher-than-expected benefit represents a gain rather than a loss. However, we observed a consistent behavioral pattern across both beneficiary-benefit and benefactor-cost manipulations, thereby effectively ruling out the “Loss Aversion” Hypothesis as an alternative explanation for the observed asymmetric adjustment.

From the perspective of the types of response indicators, to eliminate potential confounds related to introspective ability inherent in subjective emotional evaluations , and to introduce an incentivized and objective measure of participants’ responses^31,52^, we replaced the trial-by-trial subjective gratitude ratings with trial-by-trial monetary allocations in Experiment 4. The experimental procedure was identical to that of Experiment 1, with one key modification: after learning of the co-player’s decision to help, participants completed a monetary allocation task either before or after the revelation of the final benefactor-cost. In this task, they decided how many of 20 endowed tokens to allocate to the co-player, keeping the remainder for themselves. Participants were explicitly informed that these allocations would directly determine the final financial payoffs for both themselves and their respective co-players. Consequently, the number of tokens allocated to the co-player served as an objective behavioral index of their willingness to reciprocate. The results of the monetary allocations in Experiment 4 mirrored the subjective gratitude ratings from Experiment 1, with participants’ monetary allocations exhibiting an “asymmetric adjustment” in the exact same direction (Fig. 2F-G; Tables S1 and S2). This finding not only corroborates the validity of the subjective gratitude ratings employed in Experiments 1 to 3, but also demonstrates that the “asymmetric adjustment” persists across different response indicators.

From the perspective of the magnitudes of outcomes, building on Experiment 4, Experiment 5 manipulated the magnitude of the final outcomes by introducing 3-or-7 and 4-or-6 benefactor-cost conditions (each with an equiprobable 50% distribution) alongside the original 2-or-8 condition. This design ensured that the expected value during the uncertain phase remained constant across all Uncertain-to-Certain conditions, solely manipulating the magnitude of the final outcome. Both the numeric manipulation of the Uncertain-to-Certain situation (Uncertain 2-or-8 vs. 3-or-7 vs. 4-or-6) and the phase of monetary allocation (Uncertain_LowerOutcome vs. Uncertain_OutcomeUnknown vs. Uncertain_HigherOutcome) significantly impacted monetary allocations, yielding an interaction effect (*F*(1.81, 67.05) = 31.00, *p* < 0.001, 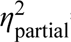 = 0.46;; Fig. 2H; Table S4). Specifically, greater deviations of the final outcome from the initial expectation (increasing from the 4-or-6, to the 3-or-7, and to the 2-or-8 conditions) elicited correspondingly larger changes in monetary allocations following the resolution of uncertainty. Despite this, when we calculated the magnitude of these changes from the uncertain to the certain phase and submitted them to an analysis of variance (ANOVA), with Numeric manipulation of the Uncertain-to-Certain situation (Uncertain 2-or-8 vs. 3-or-7 vs. 4-or-6) and Direction of change (from Uncertainty to Higher Outcome vs. from Uncertainty to Lower Outcome) as factors, we only observed two significant main effects (Numeric manipulation: *F*(1.95, 74.07) = 0.44, *p* = 0.436,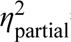 = 0.02; Direction of change: *F*(1.95, 74.07) = 0.44, *p* = 0.436, 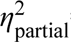 = 0.02), but no significant interaction effect (*F*(1.95, 74.07) = 0.44, *p* = 0.436, 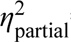 = 0.02; Fig. 2I; Table S5). This result demonstrates that although the outcome magnitude parametrically modulated the absolute degree of change in allocations before and after the resolution, the “asymmetric adjustment” robustly persists across various outcome magnitudes, and critically, its effect size is not significantly modulated by the absolute scale of the outcome deviation.

Moreover, consistent with the findings of Experiment 1, the Outcome magnitude (2 vs. 3 vs. 4 vs. 6 vs. 7 vs. 8) significantly influenced participants’ monetary allocations to the co-players (*F*(1.32, 48.67) = 105.69, *p* < 0.001,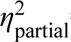 = 0.74). Though the main effect of Situation (Uncertain-to-Certain vs. Constantly-Certain) was not significant (*F*(1.00, 37.00) = 2.32, *p* = 0.136,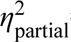 = 0.06), a significant interaction was observed between Outcome magnitude and Situation (*F*(1.59, 58.68) = 40.24, *p* < 0.001,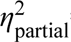 = 0.52; Fig. S1; Table S6), revealing that the main effect of the outcome was attenuated in the Uncertain-to-Certain situation than in the Constantly-Certain situation. This result provides further evidence that the transition from uncertainty to certainty involves distinct cognitive processing of outcomes compared to the Constantly-Certain baseline.

It is worth noting that, in the uncertainty phases of our experiments, we presented participants only with the potential outcomes and their associated probabilities. To further rule out the possibility that participants’ subjective expectations might deviate from theoretical mathematical expectations during the uncertainty phase^53^, in Experiments 2-7, before the interpersonal task, we asked participants to report their expectations of the average benefactor-cost (Experiments 3-7) or average beneficiary-benefit (Experiments 2–3) across the experimental conditions. We then compared these subjective expectations with the corresponding theoretical mathematical expectations. The results revealed that across Experiments 2 to 5 (as well the following Experiments 6-7), the participants’ anticipated average actual benefactor’s costs and beneficiary’s benefits did not significantly deviate from the mathematical expectations of their respective conditions (Table S7). This empirical evidence rules out subjective expectation bias as a potential confounding factor.

### Elucidating the cognitive computations of the “asymmetric adjustment”: An asymmetric intrinsic prosocial bias as the core mechanism

Having established the robust behavioral foundation of the “asymmetric adjustment” and excluded the possibilities of loss aversion or subjective expectation bias in Experiments 1-5, we advanced to the cognitive computation level to decode the latent rules generating this phenomenon. Specifically, Experiment 6 applied formal computational modeling to fit and compare competing mathematical models, aiming to dissociate whether this dynamic adjustment is governed by differential sensitivity to positive versus negative PEs or an intrinsic prosocial bias. To provide the requisite empirical variance for a rigorous model comparison, Experiment 6 utilized the 2-or-8 cost framework while systematically modulating prior probabilities; specifically, the likelihood of a high-cost outcome was varied across five levels (20%, 40%, 50%, 60%, and 80%). This parametric modulation decoupled outcome magnitudes from PE values, thereby facilitating the dissociation of distinct cognitive effects during computational modeling. Furthermore, the trial-by-trial monetary allocation, serving as an objective and incentivized behavioral index, was employed as the dependent variable, with the computational models fitted specifically to the values of reciprocal changes observed before and after the resolution of uncertainty.

Consistent with our preceding findings, model-free analyses revealed both the main effects and a significant interaction between Situation (Uncertain-to-Certain 20%8 vs. Uncertain-to-Certain 40%8 vs. Uncertain-to-Certain 50%8 vs. Uncertain-to-Certain 60%8 vs. Uncertain-to-Certain 80%8 vs. Constantly-Certain) and Phase of monetary allocation (Outcome2 vs. OutcomeUnknown/Outcome5 vs. Outcome8) on the amounts of monetary allocation (Situation: *F*(1.86, 78.04) = 66.00, *p* < 0.001,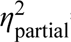 = 0.61; Outcome: *F*(1.37, 57.62) = 141.89, *p* < 0.001, 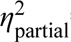 = 0.77; Interaction: *F*(2.66, 111.87) = 75.10, *p* < 0.001,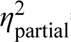 = 0.64; Fig. 3A), suggesting the differences in the magnitude of dynamic adjustment in reciprocity from exogenous uncertainty to certainty under these 5 Uncertain-to-Certain conditions, as well as the reduced sensitivity to final outcome in the Uncertain-to-Certain situation than in the Constantly-Certain situation (Table S8). Moreover, we calculated the magnitude of the changes from the uncertain to the certain phase, and estimated the effects of the computed PE values for each condition and the direction of change (from Uncertainty to Higher Outcome vs. from Uncertainty to Lower Outcome). The results showed that although the magnitude of PE significantly modulated the absolute values of changes in both the two directions (significant main effect of |PE|: *F*(2.80, 117.70) = 9.50, *p* < 0.001, 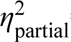 = 0.19; significant main effect of Direction of change: *F*(1.00, 42.00) = 12.03, *p* < 0.001,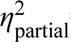= 0.22; Fig. 3B), it did not significantly influence the extent of the “asymmetric adjustment” (insignificant interaction effect: *F*(3.10, 130.33) = 0.60, *p* = 0.624,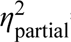= 0.01; Fig. 3B; Table S9). This provides preliminary evidence against the notion that the differential sensitivity to positive vs. negative prediction errors serves as the primary driver of the observed “asymmetric adjustment”.

**Fig. 3.**
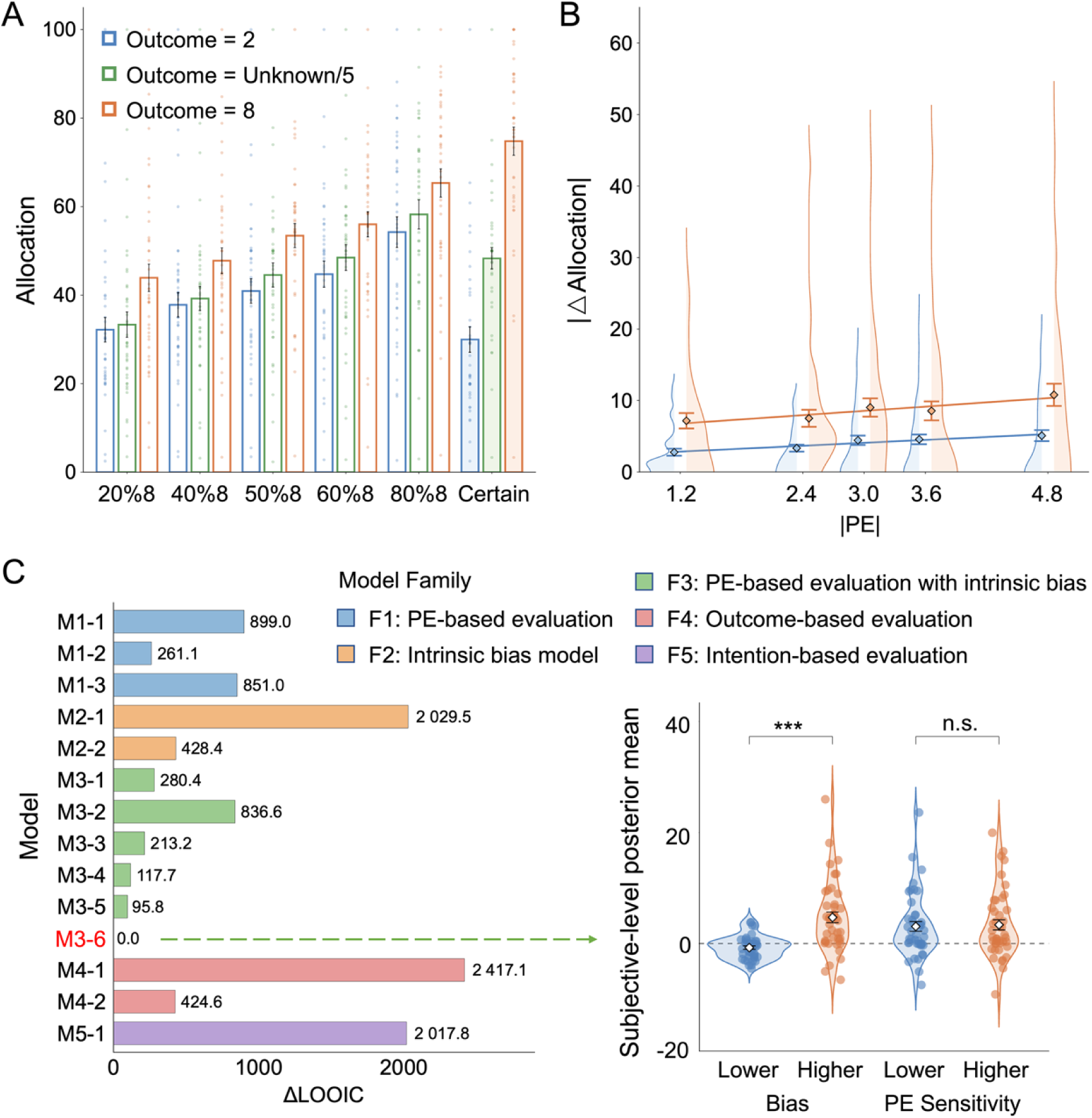
An asymmetric intrinsic prosocial bias as the core computational mechanism underlying the cognitive computations of the “asymmetric adjustment”. **(A)** Monetary allocations across conditions with systematically modulated prior probabilities (e.g., 20% to 80% chance of high outcome) and constant certainty in Experiment 6 (n = 43). **(B)** The absolute magnitude of dynamic adjustment in allocations (|△Allocation|) plotted against the absolute prediction error (|PE|). While PE magnitude modulated absolute changes, it did not significantly influence the extent of the “asymmetric adjustment”, refuting the differential PE sensitivity hypothesis. **(C)** Formal computational model comparison and parameter extraction. Model M3-6, integrating both asymmetric PE sensitivity and asymmetric baseline biases conditioned on the direction of change, provided the best fit. The subjective-level posterior means (right) reveal that the magnitude of the positive bias (|*Bias_pos_*|) when outcomes exceeded expectations was significantly larger than that of the negative bias (|*Bias_neg_*|) when outcomes fell short, whereas PE sensitivity exhibited no significant directional difference. This asymmetric intrinsic prosocial bias serves as the core cognitive mechanism driving reciprocal adjustments. Error bars represent standard errors. \*\*\**p* < 0.001, n.s. *p* > 0.05.

Through formal computational modeling, we fitted and compared 14 models that were built on the PE-based evaluation hypothesis (M1.1-M1.3), the “Intrinsic Prosocial Bias” Hypothesis (M2.1-M2.2), the combinations of these two mechanisms (M3.1-M3.6), as well as the baseline control models of outcome-based evaluation (M4.1-M4.2) and intention-based evaluation (M5.1), respectively (Table S10). Model comparison revealed that the model that incorporated both asymmetric PE sensitivity and asymmetric baseline biases conditioned on the direction of change (from Uncertainty to Higher Outcome vs. from Uncertainty to Lower Outcome) outperformed other models, with good performance of prediction (*β* = 0.94 ± 0.02, *t*(2475) = 53.94, *p* < 0.001, *r^2^*= 0.54) and parameter recovery (*r* = 0.96 ± 0.01, 95%CI = [0.94, 0.97], *t*(158) = 40.45, *p* < 0.001; Tables S10-S13; Fig. S2). The core assumption of this model was that the participant’s reciprocal adjustment, following the resolution of exogenous uncertainty, was driven not only by the PE but was also modulated by intrinsic baseline biases. The respective impact of both factors on the changes in reciprocity varies depending on the direction of the outcome shift, mathematically captured by the parameters *α_pos_*, *α_pos_*, and *Bias_pos_*, *Bias_neg_* (Eq. 1).

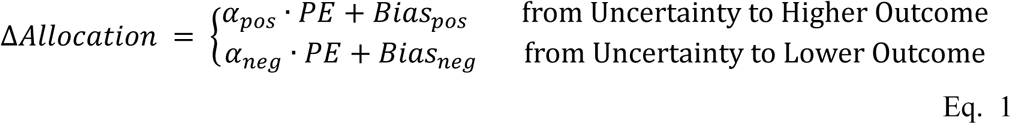

To be noted, while this combined model provided the best fit, a direct comparison of the extracted PE sensitivity parameters across the two directions of change yielded no significant difference (*t*(42) = 0.27, *p* = 0.786, Cohen’s *d* = 0.04; Fig. 3C; Tables S11 and S12). This parametric finding aligns with the model-free results, further corroborating that the “Differential PE Sensitivity” Hypothesis could not fully explain the “asymmetric adjustment”. Conversely, an analysis of the extracted bias parameters revealed an asymmetric effect (|*Bias_pos_*| vs. |*Bias_neg_*|: *t*(42) = 5.34, *p* < 0.001, Cohen’s *d* = 0.81; Fig. 3C; Tables S11 and S12): the magnitude of the positive bias when the final benefactor-cost exceeded expectations (mean = 4.70 ± 0.97, significant larger than zero: *t*(42) = 5.34, *p* < 0.001, Cohen’s *d* = 0.81) was significantly larger than that of the negative bias when the cost fell short of expectations (mean = -0.74 ± 0.31, significant lower than zero: *t*(42) = 5.34, *p* < 0.001, Cohen’s *d* = 0.81). In other words, when the benefactor’s actual cost exceeded expectations, beneficiaries applied a pronounced positive bias to reward the augmented effort; however, when the benefactor’s actual cost fell below expectations, the resulting negative bias was severely attenuated, avoiding a symmetrical, reduction in reciprocal behavior.

### Mapping the biological implementation: The exogenous uncertain-to-certain transition shifts outcome processing from the reward network toward the theory-of-mind network

Our behavioral and computational modeling findings suggest that, an asymmetric intrinsic prosocial bias, rather than an asymmetric PE-based evaluation, may be the core cognitive mechanism underlying the “asymmetric adjustment” in the beneficiary’s reciprocal behavior during the exogenous uncertain-to-certain transition. If this is the case, the beneficiary’s cognitive and neural processing upon the revelation of actual outcomes in the Uncertain-to-Certain situation should fundamentally differ from the processing of identical outcomes in the Constantly-Certain situation. Therefore, in Experiment 7, we advanced to the biological implementation level to provide neurobiological evidence supporting these computationally identified cognitive mechanisms. By integrating the behavioral paradigm from Experiment 4 with fMRI multivariate pattern analysis (MVPA), we first pinpointed the neural representations supporting the “asymmetric adjustment.” Subsequently, leveraging meta-analytical neural decoding, we mapped these identified neural patterns onto specific cognitive processes, thereby validating our cognitive computational findings.

Behaviorally, consistent with the findings from Experiments 1-4, participants in Experiment 7 robustly exhibited the “asymmetric adjustment” in their reciprocity, while also demonstrating differential sensitivity to outcomes between the Uncertain-to-Certain and Constantly-Certain situations (Fig. S3; Tables S1-S3). Consequently, we first examined the similarities and differences in the neural patterns of outcome processing between these two situations at the whole-brain level. Specifically, we employed the linear Support Vector Machine (SVM)^54,55^ to train two multivariate pattern classifiers: (1) Uncertain-to-Certain classifier, which discriminated the neural responses of Uncertain_Outcome8 and the Uncertain_Outcome2 conditions (Fig. 4A; force-choice classification *accuracy* = 58.7 ± 8.7%, *p* = 0.301, non-force-choice classification *accuracy* of 63.0 ± 6.6%, *p* = 0.016), and (2) Constantly-Certain classifier, which discriminated the neural responses of Certain_Outcome8 and the Certain_Outcome2 conditions (Fig. 4B; force-choice classification *accuracy* of 71.7 ± 10.6%, *p* = 0.005, non-force-choice classification *accuracy* of 68.5 ± 7.1%, *p* < 0.001) (see *Methods*). In order to compare the differences in neural representations between the Uncertain-to-Certain and the Constantly-Certain situations, we performed cross-situation classification, i.e., examining whether the Uncertain-to-Certain classifier can distinguish the contrast maps of the two conditions in Constantly-Certain situation and vice versa. Results of cross-situation classification revealed that the Uncertain-to-Certain classifier was able to distinguish between Certain_Outcome8 and Certain_Outcome2 conditions (force-choice classification *accuracy* = 69.6% ± 10.3%, *p* = 0.011; Fig. 4A), but the Constantly-Certain classifier was not able to distinguish between Uncertain_Outcome8 and Uncertain_Outcome2 conditions (force-choice classification *accuracy* = 63.0% ± 9.3%, *p* = 0.104; Fig. 4B). These evidence at whole-brain level indicated that, compared with the Constantly-Certain situation, there might existed unique neural representations in the Uncertain-to-Certain situation.

**Fig. 4.**
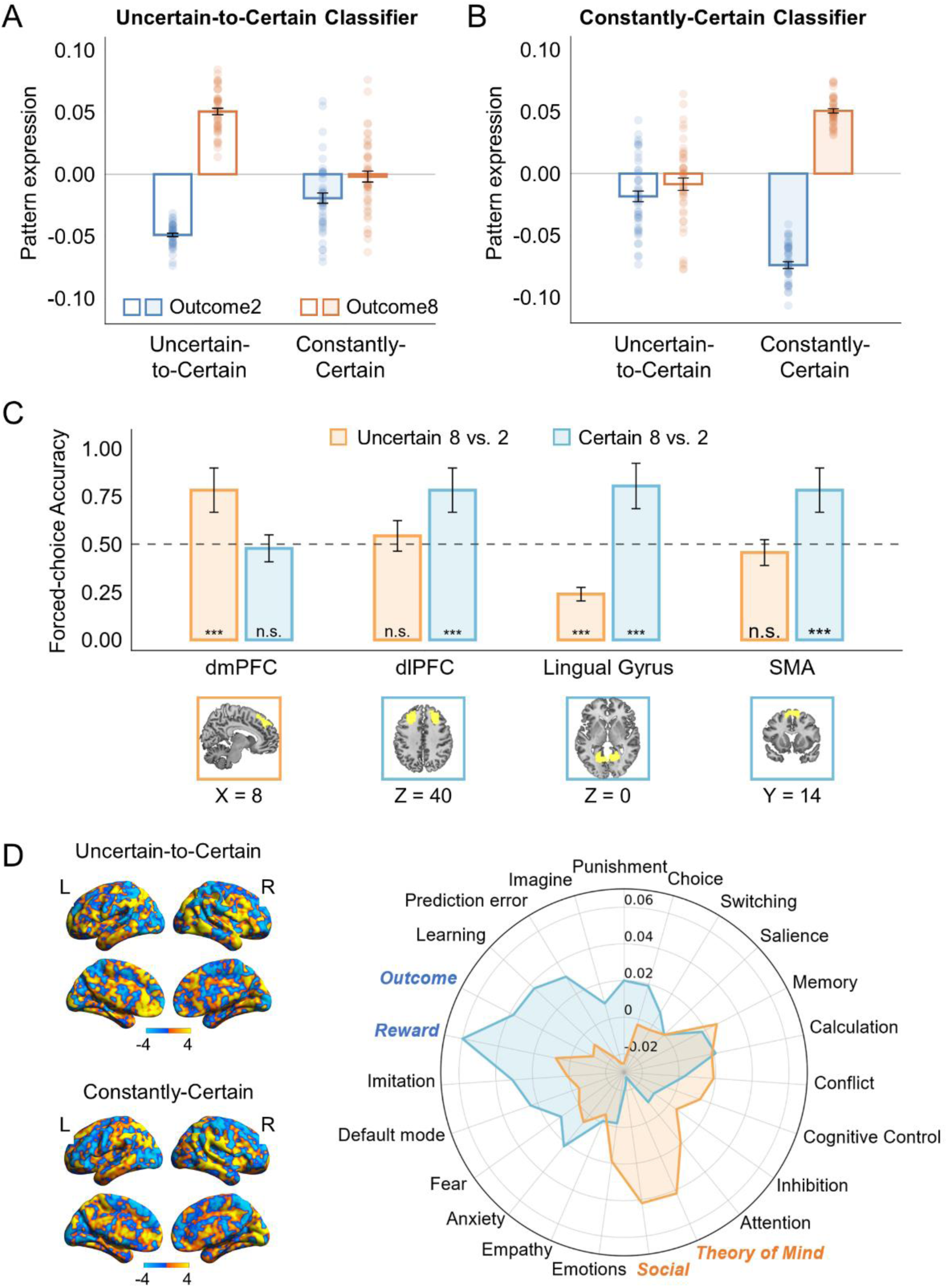
Differential neural representations for outcome processing in the Uncertain-to-Certain and in the Constantly-Certain situations. (A-B) Pattern expression values of the Uncertain-to-Certain classifier (A) and the Constantly-Certain classifier (B) in the four conditions respectively (n = 46). **(C)** Classification accuracies for Uncertain_Outcome8 vs. Uncertain_Outcome2 conditions (orange bar) and Certain_Outcome8 vs. Certain_Outcome2 conditions (blue bar) in regions that showed differential sensitivities to the Uncertain-to-Certain situation and the Constantly-Certain situation (n = 46). Results were thresholded at *p* < 0.05, Bonferroni corrected, two-tailed. **(D)** Weight maps of whole-brain multivariate pattern classifiers and Meta-analytical decoding results. The left side of the figure is the weight maps of whole-brain multivariate pattern classifiers discriminating Uncertain_Outcome8 vs. Uncertain_Outcome2 conditions (the Uncertain-to-Certain classifier) and Certain_Outcome8 vs. Certain_Outcome2 conditions (the Constantly-Certain classifier), respectively. The right side of the figure were meta-analytical decoding results. The orange line represents the similarities between the meta-analytical maps of terms of psychological components generated from Neurosynth database and the Uncertain-to-Certain classifier, and the blue line corresponds to these for the Constantly-Certain classifier. The results showed that the processing in the Uncertain-to-Certain situation was more strongly linked to “Theory of Mind” and “Social” terms, whereas that in the Constantly-Certain situation was more closely associated with “Outcome” and “Reward” terms.

To search for specific brain regions that involved in the unique neural representations in the Uncertain-to-Certain situation, we conducted ROI-based MVPA within the gratitude- and reciprocity-related brain regions identified using representational similarity analysis (RSA; see *SI Methods* and Fig. S4) and applied linear SVM^54,55^ to train local classifiers discriminating Uncertain_Outcome8 vs. Uncertain_Outcome2 conditions. This analysis revealed only one brain parcel in the dmPFC showing significant classification ability for Uncertain_Outcome2 and Uncertain_Outcome8 conditions (force-choice classification *accuracy* = 78.3% ± 11.5%, *p* < 0.001, *p*_Bonferroni_= 0.010, Fig. 4C). Meanwhile, the neural representations in these parcels could not discriminate Certain_Outcome8 vs. Certain_Outcome2 conditions (force-choice classification *accuracy* = 47.8% ± 7.1%, *p* = 0.883, *p*_Bonferroni_= 1.000). These results demonstrated that the dmPFC, a region that plays an important role in the processing of ToM as suggested by previous studies^35,56–59^, was specifically involved in the outcome processing in the Uncertain-to-Certain situation.

We conducted similar analyses as the Uncertain-to-Certain situation to search for specific brain regions involved in the Constantly-Certain situation. This analysis revealed three brain parcels that could discriminate Certain_Outcome2 and Certain_Outcome8 conditions, locating in bilateral dlPFC (force-choice classification *accuracy* = 78.3% ± 11.5%, *p* < 0.001, *p*_Bonferroni_ = 0.010), SMA (force-choice classification *accuracy* = 78.3% ± 11.5%, *p* < 0.001, *p*_Bonferroni_= 0.010), and lingual gyrus (force-choice classification *accuracy* = 80.4% ± 11.9%, *p* < 0.001, *p*_Bonferroni_= 0.003), respectively (Fig. 4C). Moreover, the neural representations in these parcels could not discriminate Uncertain_Outcome8 and Uncertain_Outcome2 conditions (dlPFC, force-choice classification *accuracy* = 54.3% ± 8.0%, *p* = 0.659, *p*_Bonferroni_= 1.000; SMA, force-choice classification *accuracy* = 45.7% ± 6.7%, *p* = 0.659, *p*_Bonferroni_= 1.000; lingual gyrus, force-choice classification *accuracy* = 23.9% ± 3.5%, lower than 50%, *p* < 0.001, *p*_Bonferroni_ = 0.035), indicating that these regions were specifically involved in the outcome processing in the Constantly-Certain situation.

Given that both evidence from the whole-brain and ROI-based analyses suggested that the outcome processing in the Uncertain-to-Certain and the Constantly-Certain situations may involve differential cognitive components, we formally test this notion by performing meta-analytic decoding for the whole-brain Uncertain-to-Certain classifier and the whole-brain Constantly-Certain classifier separately using the Neurosynth database^60^. The results showed the differences in cognitive processing between the two situations: the processing in the Uncertain-to-Certain situation was more strongly linked to “Theory of Mind” and “Social” terms, whereas that in the Constantly-Certain situation was more closely associated with “Reward”, “Outcome”, “Learning” and “Prediction Error” terms (Fig. 4D). These results profoundly enrich our mechanistic understanding of the “asymmetric adjustment”. On the one hand, the attenuated neural representation of prediction errors in the Uncertain-to-Certain situation, relative to the Constantly-Certain baseline, provides further neurobiological evidence ruling out the “Differential PE Sensitivity” Hypothesis, which further validate our computational modeling results. On the other hand, these findings highlight the theory-of-mind system as the crucial neural basis supporting the asymmetric intrinsic prosocial bias and the resultant “asymmetric adjustment” in the beneficiary’s reciprocity.

### Revealing the adaptive function of the “asymmetric adjustment”: A social signaling mechanism in sustaining cooperation

Having mapped the underlying cognitive computations and neural substrates in Experiments 6 and 7, we advanced to the adaptive functional level to address the social utility of this behavior. Crucially, these above findings highlight the theory-of-mind system as the crucial neural basis supporting the asymmetric intrinsic prosocial bias and the resultant “asymmetric adjustment” in the beneficiary’s reciprocity. The recruitment of the theory-of-mind system provides a mechanistic clue, raising the possibility that the beneficiary may infer and consider the benefactor’s reactions and the interpersonal consequences of their reciprocal behavior. Considering the adaptive role of gratitude and the resulting reciprocity^61–63^, it is plausible to suggest that this specific response pattern may help beneficiaries garner a higher level of social acceptance and reputation, thereby sustaining human cooperation when exogenous uncertainty leads to unexpected outcomes. If this is the case, then compared with alternative reciprocal strategies, the “asymmetric adjustment” should be more accepted and appreciated by other social members.

To test this hypothesis, an additional sample of participants were recruited to complete a third-party social evaluation questionnaire (Experiment 8). In the questionnaire, each participant made social evaluation as a third-party on beneficiaries, who completed an interpersonal game that was similar as previous experiments and exhibited different patterns of gratitude-induced reciprocity (Fig. 5A; see *Methods*). Here, Complete Adaptive Asymmetric Adjustment (CAAA) and Incomplete Adaptive Asymmetric Adjustment (IAAA) corresponded to the “asymmetric adjustment”, Symmetric Adjustment & Higher Amount (SAH) and Symmetric Adjustment & Lower Amount (SAL) corresponded to the pure outcome-oriented evaluation, No Adjustment (NAD) corresponded to the pure intention-oriented evaluation, and Selfish Asymmetric Adjustment (SAA) corresponded to the pattern opposite to the “asymmetric adjustment”. In line with our hypothesis, results of one-way ANOVA showed that individuals displaying “Complete Adaptive Asymmetric Adjustment” or “Incomplete Adaptive Asymmetric Adjustment” patterns were perceived as more moral (*F*(3.29, 158.03) = 33.34, *p* < 0.001), more well-meaning (*F*(3.51, 168.26) = 39.24, *p* < 0.001), more likable (*F*(3.04, 145.72) = 35.19, *p* < 0.001), less stingy (*F*(3.68, 176.51) = 34.22, *p* < 0.001) and less utilitarian (*F*(3.78, 181.39) = 20.46, *p* < 0.001) than others (Fig. 5B and Table S14), and the third-party participants were more inclined to become friends with them (*F*(3.46, 166.30) = 28.32, *p* < 0.001, Fig. 5C). Consistent with our hypothesis, these results suggest that the asymmetric intrinsic prosocial bias and the resultant “asymmetric adjustment” can indeed make the beneficiaries be perceived as morally superior, and promote their cooperative relationships with the benefactor.

**Fig. 5.**
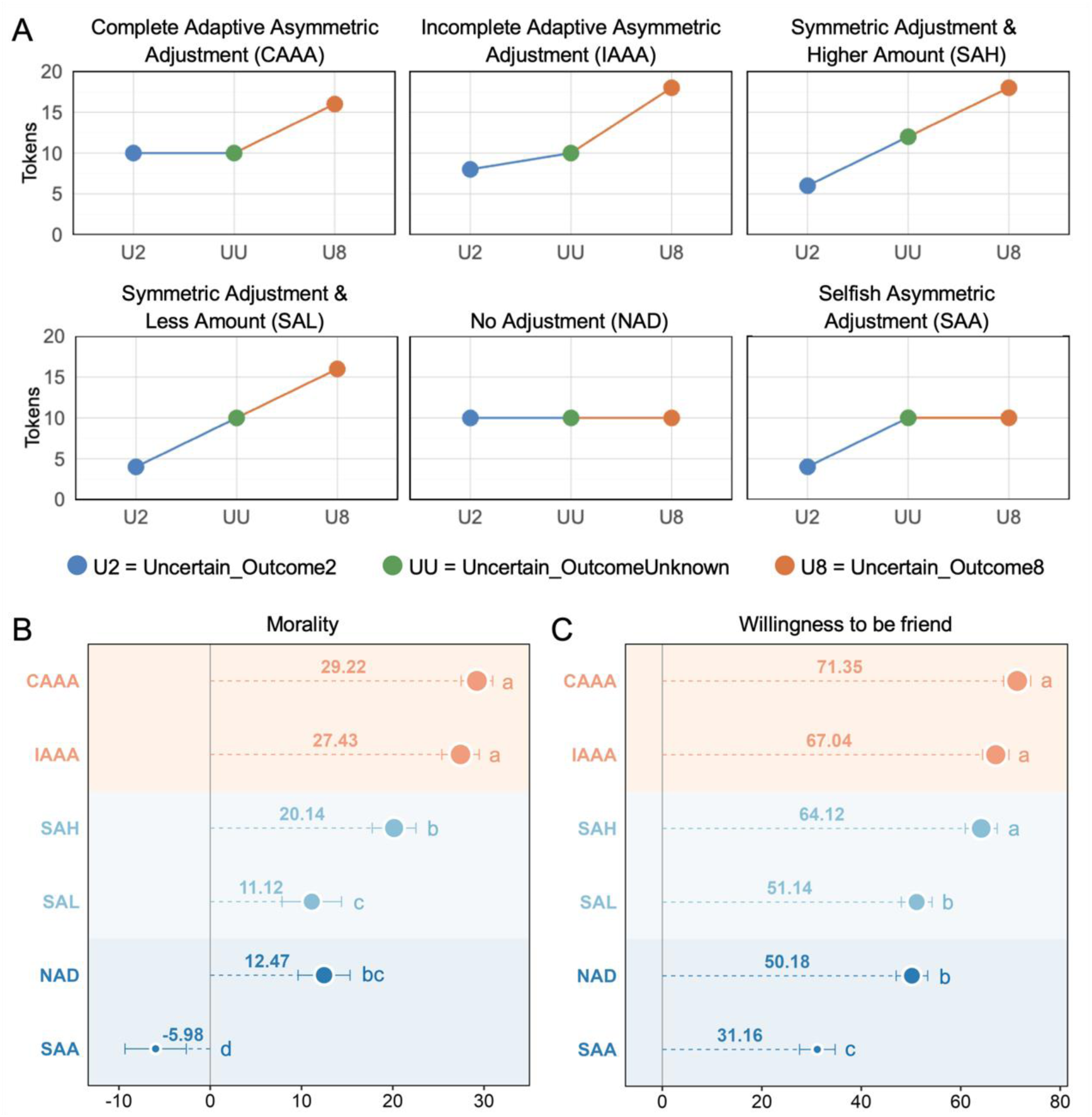
Third-party evaluations revealing the adaptive function of the “asymmetric adjustment”. **(A)** Six different reciprocal patterns in the questionnaire. Participants (n = 49) were asked to make social evaluation as a third-party on beneficiaries, who completed an interpersonal game that was similar to our fMRI experiment and exhibited different patterns of reciprocity. Here, Complete Adaptive Asymmetric Adjustment (CAAA) and Incomplete Adaptive Asymmetric Adjustment (IAAA) corresponded to the “asymmetric adjustment”, Symmetric Adjustment & Higher Amount (SAH) and Symmetric Adjustment & Lower Amount (SAL) corresponded to the pure outcome-oriented evaluation, No Adjustment (NAD) corresponded to the pure intention-oriented evaluation, and Selfish Asymmetric Adjustment (SAA) corresponded to the pattern opposite to the “asymmetric adjustment”. **(B-C)** Individuals displaying “Complete Adaptive Asymmetric Adjustment” or “Incomplete Adaptive Asymmetric Adjustment” patterns were perceived as more moral than others (B), and the third-party participants were more inclined to become friends with them (C). Different letters above the boxes indicate significant differences between groups (corrected *p* < 0.05, one-way ANOVA). Error bars represent standard errors.

## Discussion

Through eight experiments, this study provides a comprehensive, mechanistically grounded account of the “asymmetric adjustment” in the beneficiary’s affective evaluation and reciprocity during the exogenous uncertain-to-certain transition. We found that the beneficiary’s gratitude and ensuing reciprocity intensify when the final outcome (benefactor-cost or self-benefit) exceeds initial expectations; however, a parallel reduction does not elicit an equivalent decrease. By adopting a levels-of-analysis framework^50,51^, we integrated findings across three complementary dimensions to decode this phenomenon. At the cognitive computation level (Marr’s algorithmic level), computational modeling demonstrates that this dynamic adjustment is driven by an asymmetric intrinsic prosocial bias integrated into the valuation process. At the biological implementation level (Marr’s implementational level), our findings highlight the theory-of-mind system, particularly the dorsomedial prefrontal cortex, as the neural basis supporting these dynamic computations. Finally, at the adaptive social function level (Marr’s computational level), third-party evaluations reveal that this bias is perceived as morally superior to alternative reciprocal strategies. Together, these multi-level findings offer new insights for the computational resilience of human cooperation: by amplifying unexpected altruistic signals while buffering against shortfalls, the observed cognitive asymmetry serves an essential adaptive function in in sustaining human cooperation when environmental “noises” introduce unavoidable discrepancies between expectation and reality.

The observed “asymmetric adjustment” challenges existing theoretical frameworks at multiple levels. Within the specific domain of gratitude and reciprocity, classical accounts, whether focused on outcome-based evaluation^3,28,35,37,42^, intention-based evaluation^3,28,34,37,38,41^, symmetrical PE-based social learning^13^ or their linear integration, consistently predict symmetric rather than asymmetric adjustments in exogenous uncertainty-to-certainty transitions. Beyond these domain-specific theories, our findings further exclude other explanations from broader domains. First, by manipulating both benefactor-cost and beneficiary-benefit (Experiments 2-3), we demonstrate that the observed “asymmetric adjustment” could not be fully explained by the tendency of loss aversion proposed by the Prospect Theory^43–45^. Second, by comparing subjective ratings with the corresponding theoretical values (Experiments 2-7), we further rule out the possibility that participants’ subjective expectations might deviate from theoretical mathematical expectations during the uncertainty phase^53^. Third, through computational modeling (Experiment 6) as well as the meta-analytical neural decoding (Experiment 7), we show that the phenomenon cannot be fully accounted for by the differential sensitivity to positive and negative prediction errors (PEs)^46–48^. Instead, our winning model identifies an asymmetric intrinsic prosocial bias as the core mechanism. This bias is manifested as a positive baseline offset (intercept) when the final benefactor-cost exceeds expectations, but an insignificant offset when the cost falls short of expectations. This suggests that beneficiaries navigating exogenous uncertainty are not “biased evaluators” of prediction errors, but rather “outcome-dependent generous evaluators” from the outset.

From an evolutionary perspective, the mechanism of asymmetric intrinsic prosocial bias identified here aligns with the Error Management Theory (EMT)^64^. This theory posits that cognitive biases in perception, attention, and social judgments can be understood as the products of evolutionary section; whenever the costs of false positive and false negative errors are asymmetric under uncertainty, selection would favor a cognitive bias toward the least costly error. Our findings suggest that in the context of reciprocity under exogenous uncertainty, this cognitive bias may be outcome-dependent. For one thing, when a benefactor’s actual cost exceeds expectations, failing to consolidate a highly cooperative partner, a false negative incurred by reducing or merely maintaining the level of reciprocity established during the uncertainty phase, carries a high evolutionary cost, as it leads to the permanent loss of a valuable cooperative relationship. Conversely, overestimating their goodwill (a false positive) only has a short-term impact due to a single over-repayment. Consequently, selection favors the positive bias we observed, which acts as a strategic amplifier to rapidly lock in high-quality partners.

For another, when a benefactor’s actual cost falls short of expectations, a beneficiary whose reciprocity is strictly determined by objective outcomes would be expected to exhibit a negative bias equivalent in magnitude to the positive bias observed when costs exceed expectations. However, our results demonstrate a significant asymmetry: while a negative bias is present, its magnitude is severely attenuated compared to the robust positive bias. Error Management Theory^64^ provides a functional explanation for this asymmetric scaling. In contexts of exogenous uncertainty, unfavorable outcomes frequently result from environmental noise rather than uncooperative intentions. Imposing a proportionally large negative bias would substantially increase the risk of a false-negative error, specifically, permanently terminating a relationship with a high-quality partner whose efforts were merely diminished by external factors. By sharply restricting the magnitude of the negative bias, beneficiaries effectively buffer against this catastrophic social loss. At the same time, maintaining a slight negative bias, rather than displaying a completely zero or positive bias in unfavorable situations, prevents extreme false-positive errors, thereby protecting the beneficiary from persistent exploitation by genuine free-riders. Therefore, in this context, this heavily attenuated negative bias confers the greatest social adaptive advantage, balancing the vital need for relationship preservation with basic self-protection. Viewed through these lenses, the observed “asymmetric adjustment” and its underlying mechanism of an asymmetric intrinsic prosocial bias may serve to minimize the overall cost of social navigation, playing a pivotal role in sustaining interpersonal cooperation. Indeed, this is supported by our third-party evaluation evidence (Experiment 8), showing that individuals exhibiting such asymmetric adjustment are perceived as more morally superior, suggesting this computational strategy is heavily favored by social selection.

The mechanism of asymmetric intrinsic prosocial bias aligns with previously documented phenomena in broader social cognition. Our finding, specifically, the presence of a robust positive bias during favorable outcomes and the suppression of a negative bias during unfavorable ones, echoes the well-established person-positivity bias^65^ and positive illusions in close relationships^66^. These phenomena collectively demonstrate a systematic cognitive preference to view interacting partners favorably to sustain social bonds. By employing formal computational modeling, our study not only allows the extension of Error Management Theory^64^ to the dynamic computation of reciprocity under exogenous uncertainty, but also offers critical cognitive-computational insights for understanding other adaptive cognitive biases.

At the neural level, MVPA and neural decoding revealed that under uncertain-to-certain conditions, the processing difference between outcomes that were higher or lower than expected primarily involved theory-of-mind-related process, notably in the dmPFC^20,30,67^. In contrast, under certain conditions, the corresponding outcome differences were mainly associated with outcome and reward-related processing, involving regions such as the dlPFC^20,30,68^. These findings indicate that despite identical final outcomes, the transition from uncertainty to certainty shifts the brain’s final outcome processing from a value-based computation toward a mental-state inference process. This process may play a pivotal role in anticipating the benefactor’s reactions and the interpersonal consequences elicited by one’s own reciprocal behavior across varying direction of outcome transitions, and integrating these social considerations with broader reputational and cooperative goals. This finding aligns with prior research^12,69–72^, underscoring the critical role of theory-of-mind system in navigating interpersonal processes under uncertainty.

This study provides important implications for future research and daily life decision-making. First, our study identifies a novel phenomenon of adaptive asymmetric adjustment in gratitude and reciprocity, which is supported by an asymmetric intrinsic prosocial bias in the valuation process. This mechanism diverges from expectation-violation-based social learning frameworks previously proposed for endogenous uncertainty processing^7^, suggesting potential differential neurocognitive bases for processing exogenous versus endogenous uncertainty. While our design did not directly compare these uncertainty types, this distinction represents a critical hypothesis for future research. Moreover, our focus on one-shot exogenous uncertainty-to-transitions raises further questions about how beneficiaries manage repeated exogenous uncertainty transitions. Specifically, whether such processes engage recursive social learning mechanisms and how they contrast with the learning architectures adapted to endogenous uncertainty remain open questions warranting systematic investigation. These unresolved issues highlight promising avenues for advancing our understanding of exogenous uncertainty processing in prosocial decision-making.

Second, the identification of adaptive asymmetric adjustment in reciprocity has profound implications for computational psychiatry and artificial intelligence. In computational psychiatry, the dysregulation of this adaptive mechanism may provide a novel, quantifiable phenotype for understanding social dysfunctions. For instance, rather than exhibiting asymmetric intrinsic prosocial bias, individuals with interpersonal dysfunctions might exhibit rigid or maladaptive updating: they may fail to mount the positive bias needed to consolidate social bonds during favorable outcomes, or conversely, inappropriately apply a negative bias during noisy, unfavorable outcomes, thereby impairing the maintenance of social relationships. For example, studies utilizing iterative Trust Games have revealed that when cooperation ruptures (e.g., after receiving a low, uncooperative offer), healthy individuals typically employ “coaxing” behaviors, making unusually generous offers to signal appeasement and restore the social bond. In contrast, Borderline Personality Disorder (BPD) patients consistently fail to exhibit these reparative, generous gestures following social ruptures, leading to the irretrievable breakdown of trust^11,73^. Consequently, the specific intercept parameters established in our asymmetric model could serve as precise computational markers for assessing individual social resilience. Beyond human interaction, these computational principles may help to revolutionize human-computer (AI) interaction (HCI). If artificial intelligence systems were designed to emulate this asymmetric cooperative resilience, such as rewarding unexpected human goodwill, while adopting a cautious, non-punitive baseline (rather than strictly penalizing errors) when humans fall short due to exogenous noise, it would significantly enhance mutual trust and the long-term sustainability of human-computer (AI) cooperation^74^.

In summary, the present study uncovers a robust “asymmetric adjustment” in the beneficiary’s reciprocity under exogenous uncertainty, demonstrating a computational resilience that sustains human cooperation despite environmental noise. This adjustment transcends traditional models of outcome/intention-based calculations, loss aversion, or prediction-error-based social learning, being primarily governed by an asymmetric intrinsic prosocial bias that amplifies reciprocity for positive outcome deviations while buffering against shortfalls. At the neural level, this process is supported by representations within the theory-of-mind system, particularly the dorsomedial prefrontal cortex. Third-party evaluations further reveal that this bias is perceived as morally superior to alternative strategies, highlighting its critical adaptive function in maintaining social bonds amidst the volatility of an uncertain world. By integrating cognitive, neural, and social functional levels of analysis, our findings offer a comprehensive mechanistic account of how human reciprocity remains resilient in the face of environmental unpredictability.

## Methods

### Participants

To systematically investigate the dynamic adjustment of reciprocity under exogenous uncertainty, we adopted an integrated levels-of-analysis framework across eight experiments. These experiments were designed to delineate the phenomenon at three complementary dimensions: its fundamental behavioral manifestation (Experiments 1–5), its governing cognitive computational rules (Experiment 6), its biological implementation in the brain (Experiment 7), and its adaptive social function (Experiment 8). Undergraduate and graduate students were recruited from Shanghai, China, or via a Chinese online experiment platform (NAODAO: www.naodao.com)^75^. After excluding a minor portion of participants due to failing comprehension tests (in total 18 in Experiments 2, 3, 5, and 6) or excessive head motion (3 in Experiment 7, fMRI experiment), the final sample sizes for each experiment were: Experiment 1 (n = 33; 23 females; 21.85 ± 1.64 years), Experiment 2 (n = 31; 22 females; 21.4 ± 1.6 years), Experiment 3 (n = 33; 16 females; 21.4 ± 2.0 years), Experiment 4 (n = 37; 24 females; 20.62 ± 1.66 years), Experiment 5 (n = 38; 25 females; 20.45 ± 2.58 years), Experiment 6 (n = 43; 28 females; 20.95 ± 2.34 years), Experiment 7 (n = 46; 23 females; 21.7 ± 2.1 years), and Experiment 8 (n = 49; 21 females; 21.9 ± 2.0 years). For the fMRI study (Experiment 7, fMRI experiment), all participants were right-handed with normal or corrected-to-normal vision. None of the participants reported any history of psychiatric, neurological, or cognitive disorders. All experiments were carried out in accordance with the Declaration of Helsinki and were approved by the Ethics Committee of East China Normal University. Informed written consent was obtained from each participant prior to participating. See details in *SI Methods*.

### Procedures

#### Overview

Apart from Experiment 8 (online questionnaire), all the experiments mainly consisted of two sessions. In the first session (noise/pain titration), we measured each participant’s noise/pain threshold and determined the intensity of noise/pain was what the participant considered to be a moderate punishment. In the second session (main task), the participants (the beneficiaries) performed an interpersonal task, in each round of which they would receive a noise/pain stimulation and were randomly paired with an anonymous co-player (the benefactor), who could decide whether to help them reduce the number of noises/pains by undertaking a number of noises/pains. In Experiments 2-7, we additional included a subjective expectations rating session before the interpersonal task, where we asked participants to report their expectations of the average benefactor-cost (Experiments 3–7) or average beneficiary-benefit (Experiments 2–3) across the experimental conditions.

#### Noise/Pain titration

In Experiments 1-4, noise stimulation was employed as the punishment, a choice driven by its ease of administration and suitability for the online experiment (Experiment 3). However, in Experiments 5-8, we transitioned to pain stimulation. This adjustment was made to avoid potential confounds with the ambient acoustic noise generated by the MRI scanner, thereby ensuring the saliency and integrity of the experimental punishment. The two punitive stimuli have exactly the same effect in creating the help-receiving situation.

In order to conduct the noise titration, all the participants were asked to wear headphones and hear a clip of metal-cutting noise. We gradually increased the volume of this clip of noise until the participant reported 8 on a 10-point pain scale (1 = not noisy, 10 = intolerable), which was a moderate punishment for the participant.

Pain titration was conducted following the procedure outlined in previous studies on gratitude^34,35,41^. During the titration process, an intra-epidermal needle electrode was attached to the back of each participant’s left hand for cutaneous electrical stimulation^76^. The initial pain stimulation consisted of eight repeated pulses, each with an intensity of 0.2 mA and a duration of 0.5-ms, with a 10-ms interval between each pulse. We progressively increased the intensity of each single pulse until the participant rated the pain as 8 on a 10-level pain scale (1 = not painful, 10 = intolerable).

All participants were informed that the noise/pain stimulation that each individual would receive in the interpersonal task would be the one that they rated as 8 in the noise/pain titration session. This setting made participants aware that the intensity of the noise/pain subjectively experienced by each individual in the interpersonal interaction task was the same.

#### Subjective Expectations Rating

To further rule out the possibility that participants’ subjective expectations might deviate from theoretical mathematical expectations during the uncertainty phase, in Experiments 2-7, we asked participants to provide subjective estimates prior to the interpersonal task. Specifically, participants reported their expectations of the average benefactor-cost (in Experiments 3-7) or average beneficiary-benefit (in Experiments 2-3) they expected to occur across the respective experimental conditions. For example, to assess expectations regarding the average benefactor-cost, participants answered the following question: “Assume that the co-player chooses to offer help across multiple rounds where the cost is either 2 or 8 pain stimuli with unknown probabilities. Please estimate the average number of pain stimuli actually borne by the co-player in these rounds.” Participants provided their estimates on a 9-point scale (1 = far fewer than 5 times, 9 = far more than 5 times). These subjective estimates were subsequently compared with theoretical expected values of the experimental design to ensure no subjective expectation bias existed.

#### Interpersonal task (main task)

##### Experiment 1—Behavioral experiment manipulating uncertainty in benefactor’s cost

As the essential prerequisite for our multi-level inquiry, Experiments 1-5 were conducted to establish the empirical foundation and characterize the robustness of the ‘asymmetric adjustment’ across varied behavioral and outcome contexts.

In the Experiment 1, we used a multi-round interpersonal task (Fig. 1B) to induce dynamic changes from uncertainty to certainty in help-receiving situation, which was adapted from the previous study on how the uncertainty in benefactor’s cost influenced beneficiary’s gratitude^41^. Following the noise titration session, each participant was instructed on the general rules of the interpersonal task. In each trial, the participant was paired with an anonymous same-gender co-player, who was distinct from the ones in any other trials and would only interact with the participant once. In each round, the participant was to receive 10 times of noise stimulation of Level 8. The participant was instructed that each co-player had come to the lab before the participant and had already decided whether or not to help the participant reduce half the number of noise stimulation by enduring a number of noise stimulation (benefactor’s cost). Once being helped, the number of noise stimulation that the participant had to endure would be reduced to 5 times, regardless of the co-player’s cost; otherwise, the noise stimulation would continue to be 10 times. The participant was informed that each co-player decided whether to help the participant under one of the two situations regarding the cost of help, the Uncertain-to-Certain situation and the Constantly-Certain situation, which was randomly chosen by the computer program.

In each trial of the Uncertain-to-Certain situation, the co-player decided whether to help the participant under exogenous uncertain cost of receiving the noise stimulation of either 2 times or 8 times, each with 50% probability, i.e., the mathematic expectation of final outcome equaled 5, determined by the computer system^41^. After the decision had been made by the co-player, it cannot be changed no matter what the actual cost was. Then the actual cost that the co-player undertook, either 2 times (lower than expectation) or 8 times (higher than expectation), was determined randomly by the computer and shown to both the participant and the co-player. If the co-player offered help in this trial, the participants were instructed to rate their feelings of gratitude towards the co-player. In some of the trials of Uncertain-to-Certain situation, the participants should rate their feelings of gratitude to the co-player’s help (Response phase; 0 = not at all, 100 = very strong), at the time point before the co-player’s actual cost was presented, i.e., under uncertainty (Uncertain_OutcomeUnknown condition). In the other trials, they should rate their feeling of gratitude at the time point after knowing the co-player’s actual cost of 2 times (Uncertain_Outcome2 condition) or 8 times (Uncertain_Outcome8 condition). These three conditions in the Uncertain-to-Certain situation formed a one-way three-level within-subject design. The comparisons in the amount of self-reported gratitude rating between Uncertain_OutcomeUnknown and Uncertain_Outcome2 conditions, as well as between Uncertain_OutcomeUnknown and Uncertain_Outcome8 conditions, would reveal how the beneficiary dynamically adjusts their feeling of gratitude when transitioning from facing an uncertain outcome to facing an actual and certain outcome of higher or lower benefactor’s cost.

In each trial of the Constantly-Certain situation, the co-player decided whether to help the participant under certain cost, which could be 2, 5 or 8 times, determined randomly by the computer and displayed to the co-player prior to their decisions. Being presented with the co-player decision on whether to help or not under certain cost of 2 times (Certain_Outcome2 condition), 5 times (Certain_Outcome5 condition) or 8 times (Certain_Outcome8 condition), the participants should rate their feeling of gratitude to the co-player in the corresponding trial. The Certain_Outcome2 and Certain_Outcome8 conditions in the Constantly-Certain situation were included as baselines, along with the Uncertain_Outcome2 and Uncertain_Outcome8 conditions, forming a 2 Situation (Uncertain-to-Certain vs. Constantly-Certain) × 2 Outcome (2 vs. 8) within-subject design. This design enabled us to elucidate distinctions in the processing of gratitude across two distinct scenarios: one in which the outcome of help is ascertained from the outset, and another wherein the outcome evolves from an initial state of uncertainty to eventual certainty, despite the ultimate outcomes of benefactor’s cost being equivalent in both conditions.

The participant was also informed that, after the whole experiment, 20 trials would be randomly selected from all the trials and be realized to determine their and the corresponding co-player’s final amount of noise stimulation.

In each trial (Fig. 1B), after being paired with an anonymous co-player (Regrouping phase, 2 s), the participant would see information regarding which situation (Uncertain-to-Certain or Constantly-Certain) the co-player’s cost belonged to in the current trial (Situation information phase, 3 s). For the Uncertain-to-Certain situation, the Situation information phase would present an orange pie, with the numbers 2 and 8 on the top and the bottom, respectively (locations counterbalanced across trials), indicating the co-player’s uncertain cost of either 2 times or 8 times of pain stimulation, each with 50% probability. Then the co-player’s decision on whether to help the participant under uncertainty would display under the orange pie (Outcome_Unknown phase for Uncertain-to-Certain situation, 3s). In trials of the Uncertain_OutcomeUnknown condition (Fig. 1B), the participant should immediately rate their feelings of gratitude to the co-player paired after the Outcome_Unknown phase without knowing the co-player’s actual cost. Then the participant would see the actual outcome of co-player’s cost (Outcome_Display phase, 3s). In trials of the Uncertain_Outcome2 and Uncertain_Outcome8 conditions, the participant would see the actual outcome of co-player’s cost first (Outcome-Display phase, 3 s), and then rate their feelings of gratitude to the co-player (Response phase, < 12 s). For the Constantly-Certain situation (Fig. 1C), this phase would present an orange pie, with the number of co-player’s certain cost (2, 5 or 8) in the center. Then after the co-player’s decision on whether or not to help the participant would display under the orange pie. To parallel the two-stage rating structure of the Uncertain-to-Certain situation, the co-player’s cost was presented a second time (Outcome_Display phase, 3s), and the participant was instructed to rate their feelings of gratitude to the co-player either before or after this display (Response phase, < 12 s).

In Experiment 1, for trials in which the co-player offered help (Help trials), the Uncertain-to-Certain situation included 6 rating trials each for the Uncertain_Outcome2 and Uncertain_Outcome8 conditions, and 12 rating trials for the Uncertain_OutcomeUnknown condition. This design ensured an equal probability of eliciting gratitude ratings before versus after the revelation of the final outcome. In the Constantly-Certain situation, 6 rating trials were administered both before and after the outcome display for each of the Certain_Outcome2, Certain_Outcome5, and Certain_Outcome8 conditions. Since in the Certain situation, the participants’ ratings before and after the Outcome_Display phase were based on the same experimental information, these data were collapsed for subsequent analyses. Furthermore, we included trials in which the co-player did not offer help (30 NoHelp trials), constituting half the proportion of the Help trials (60 Help trials), to serve as fillers, yielding a total of 90 trials. The task was divided into 3 runs with equal number of trials for each condition in each run. Each run consisted of 30 trials in total and lasted for about 10 mins. Trials within each run were pseudo-randomly mixed to ensure that no more than two consecutive trials were from the same condition. To avoid the influence of trial sequence, 3 sequences with pseudo-random order of trials were pre-determined and counterbalanced across participants by using Latin square design. Unbeknown to the participants, all the co-players’ decisions were predetermined by a computer program. No participant disputed the authenticity of the co-players when reporting their comments and feelings after the experiment.

##### Experiment 2—Behavioral experiment manipulating uncertainty in beneficiary’s benefit

Experiment 2 was conducted to verify whether the “asymmetric adjustment” exists under different types of help outcomes and to test the “Loss Aversion” Hypothesis. The procedure of this behavioral experiment was the same as Experiment 1, except that (1) Experiment 2 manipulated the exogenous uncertainty in the participants’ own benefits, instead of the benefactor’s cost, and (2) to alleviate the cognitive load and potential response fatigue induced by repeated evaluations of identical information, for the Constantly-Certain situation, we only asked participants to make gratitude ratings once after the co-player’s decision and cost was presented. See trial setting in Table S15.

##### Experiment 3—Behavioral replications manipulating uncertainties in benefactor’s cost and beneficiary’s benefit simultaneously

Experiment 3 was conducted to further examine whether a consistent pattern persists across exogenous uncertainties in both the benefactor’s cost and the beneficiary’s benefit. The procedure of this behavioral experiment was the same as Experiment 2, except that the experiment included two blocks, manipulating the exogenous uncertainties in the benefactor’s cost and in the participants’ own benefits respectively. See trial setting in Table S15.

##### Experiment 4—Behavioral experiment extending the asymmetric adjustment from gratitude to gratitude-reduced reciprocity

In Experiment 4, to address potential methodological confounds related to introspective ability inherent in subjective emotional evaluations, and to introduce an incentivized, objective measure of participants’ responses, we replaced the trial-by-trial subjective gratitude ratings with actual monetary allocations to be realized after the experiment (an indicator of reciprocal behavior).

The procedure of Experiment 4 was the same as the Experiment 1, except that (1) In Experiment 4, at the Response phase, the participants were instructed to complete monetary allocation rather than rating the feelings of gratitude. Participants should allocate 20 points between themselves and the co-player in the corresponding trial (1 point = 1 Yuan, 20 Yuan ∼ 2.76 USD; the participant could adjust the amount of allocation in increments of 1 point). Meanwhile, unlike in the gratitude rating experiments, the participants were required to allocate the money in every trial, regardless of whether their co-players provided help or not. The participant was informed that all co-players were unaware of the procedure of monetary allocation, eliminating the possibility that the co-players’ decisions to help were due to monetary concerns. The participant was also informed that, after the whole experiment, 20 trials would be randomly selected from all the trials and be realized to determine the corresponding co-player’s final amount of pain stimulation and monetary bonus. The participant would receive the average amount of points that each participant assigned to themself and the average number of pain stimulation throughout the randomly selected trials. (2) Similar as Experiments 2-3, to alleviate the cognitive load and potential response fatigue induced by repeated evaluations of identical information, for the Constantly-Certain situation, we only asked participants to make monetary allocations once after the co-player’s decision and cost was presented. See trial setting in Table S15.

##### Experiment 5—Behavioral validations of the asymmetric adjustment across diverse outcome magnitude

Experiment 5 aimed to examine whether the “asymmetric adjustment” persists across diverse outcome magnitudes. To achieve this, Experiment 5 manipulated the magnitude of the final outcomes by introducing 3-or-7 and 4-or-6 benefactor-cost conditions while ensuring that the expected value during the uncertain phase remained unchanged.

The procedure of this behavioral experiment was the same as the Experiment 4, except that within the Uncertain-to-Certain situation, in addition to the 2-or-8 outcome pair (each with a 50% probability), we incorporated 3-or-7 and 4-or-6 outcome pairs (each with a 50% probability). Simultaneously, the Constantly-Certain situation encompassed all possible outcome magnitudes, namely 2, 3, 4, 5, 6, 7, and 8. Therefore, Experiment 6 comprised 16 experimental conditions: 9 Uncertain-to-Certain situation conditions, i.e., 3 (Uncertain-to-Certain Situation: Uncertain28 vs. Uncertain37 vs. Uncertain46) × 3 (Outcome in the Uncertain-to-Certain situation: Uncertain_LowerOutcome vs. Uncertain_OutcomeUnknown vs. Uncertain_HigherOutcome), and 7 Constantly-Certain situation conditions i.e., Certain_Outcome2, Certain_Outcome3, …, and Certain_Outcome8. The 9 Uncertain-to-Certain situation conditions formed a 3 (Uncertain-to-Certain Situation: Uncertain28 vs. Uncertain37 vs. Uncertain46) × 3 (Outcome in the Uncertain-to-Certain situation: Uncertain_LowerOutcome vs. Uncertain_OutcomeUnknown vs. Uncertain_HigherOutcome) within-subjects design. Each experimental condition consisted of 6 Help trials and 3 NoHelp trials, with a total of 171 trials, divided into three blocks. The block order was counterbalanced across participants using a Latin square design. See trial setting in Table S16.

##### Experiment 6—Behavioral experiment and computational modeling reveling the cognitive mechanism of “asymmetric adjustment”

Building upon the established behavioral foundation, Experiment 6 sought to decode the latent rules generating the “asymmetric adjustment.” By systematically modulating prior probabilities to decouple outcome magnitudes from prediction errors, we applied formal computational modeling to test whether this dynamic adjustment is governed by differential PE sensitivity or an intrinsic prosocial bias.

The procedure of Experiment 6 was the same as Experiment 4, except that Experiment 6 manipulated the probability of higher (i.e., 8) and lower (i.e., 2) benefactor’s cost upon the foundation of Experiment 4. Within the Uncertain-to-Certain situation, the probabilities of Outcome 2 and Outcome 8 were not limited to the 50%/50%, but also included 20%/80%, 40%/60%, 60%/40%, and 80%/20%. For brevity, these conditions were designated as the Uncertain-to-Certain 20%8, 40%8, 50%8, 60%8, and 80%8 conditions, respectively. It should be noted that to equate the number of trials for higher and lower outcomes across all probability conditions, regardless of the probability information presented during the Outcome-Unknown phase, the probabilities of Outcome 2 and Outcome 8 ultimately presented to participants were consistently 50%/50%. To prevent this setup from biasing participants’ subjective expectations of the probabilities, participants were informed that the probability information in each trial reflected the authentic probabilities considered by each co-player during their decision-making, but the outcomes the participants observed in the experiment were randomly presented after being selected by the experimenter. Therefore, Experiment 6 comprised 18 experimental conditions: 15 Uncertain-to-Certain situation conditions, i.e., 5 (Uncertain-to-Certain situation: Uncertain-to-Certain 20%8 vs. Uncertain-to-Certain 40%8 vs. Uncertain-to-Certain 50%8 vs. Uncertain-to-Certain 60%8 vs. Uncertain-to-Certain 80%8) × 3 (Outcome in the Uncertain-to-Certain situation: Uncertain_Outcome2 vs. Uncertain_OutcomeUnknown vs. Uncertain_Outcome8), and 3 Constantly-Certain situation conditions, i.e., Certain_Outcome2, Certain_Outcome5, and Certain_Outcome8. Each experimental condition consisted of 6 Help trials and 2 NoHelp trials, with a total of 144 trials, divided into three blocks. The block order was counterbalanced across participants using a Latin square design. See trial setting in Table S16.

##### Experiment 7—FMRI experiment investigating the neural bases of the “asymmetric adjustment”

Advancing to the biological implementation level, Experiment 7 aimed to provide neurobiological evidence supporting the computationally identified mechanisms in Experiment 6. The participants completed the interpersonal task in an MRI scanner.

The experimental paradigm of Experiment 7 is basically the same as that of Experiment 4, except that (1) the noise stimulus was replaced by pain stimulus; (2) to maintain the stability of the BOLD signal, the number of experimental trials in Experiment 4 is doubled; (3) for the purpose of fMRI signal deconvolution, before and after the Allocation phase and the Outcome_Display phase, a fixation was presented for 1 - 5 s. Meanwhile, the during time for the Regrouping phase and Situation information phase were set to 2 to 4 seconds. There were 12 Help trials and 6 NoHelp trials each for the conditions of Uncertain_Outcome2, Uncertain_Outcome8, Certain_Outcome2, Certain_Outcome5, and Certain_Outcome8, as well as 24 Help trials and 12 NoHelp trials for the Uncertain_OutcomeUnknown condition, ensuring that the Allocation phase occurred equally before and after the outcome of the uncertain situation was revealed. The task was divided into 3 runs, with an equal number of trials for each condition in each run. Each run consisted of a total of 42 trials and lasted for approximately 15 minutes. See trial setting in Table S15.

##### Experiment 8—Third-party evaluations revealing the adaptive function of the “asymmetric adjustment”

To further test the hypothesis regarding the adaptive function of “asymmetric adjustment” in human cooperation derived from the previous mechanistic findings, Experiment 8 recruited an additional sample of participants were to complete a third-party social evaluation questionnaire. In the questionnaire, each participant made social evaluation as a third-party on beneficiaries, who completed an interpersonal game that was similar as our fMRI experiment and exhibited different patterns of gratitude-induced reciprocity:

1. Complete Adaptive Asymmetric Adjustment (CAAA): reciprocating significantly more in Uncertain_Outcome8 (i.e., 16 points) than in Uncertain_OutcomeUnknown (i.e., 10 points), and the same as Uncertain_OutcomeUnknown in Uncertain_Outcome2 (i.e., 10 points).
2. Incomplete Adaptive Asymmetric Adjustment (IAAA): reciprocating significantly more in Uncertain_Outcome8 (i.e., 18 points) than in Uncertain_OutcomeUnknown (i.e., 10 points), and slightly less in Uncertain_Outcome2 (i.e., 8 points) than in Uncertain_OutcomeUnknown (i.e., 10 points). This pattern of reciprocity was similar to the pattern of participants’ average amounts of allocation in Experiment 4 and 7.
3. Symmetric Adjustment & Higher Amount (SAH): compared with Uncertain_OutcomeUnknown (i.e., 12 points), reciprocating more in Uncertain_Outcome8 (i.e., 18 points) and equivalently less in Uncertain_Outcome2 (i.e., 6 points), with more amount of allocation in Uncertain_OutcomeUnknown but the same total amount of allocation compared with Complete Adaptive Asymmetric Adjustment and Incomplete Adaptive Asymmetric Adjustment.
4. Symmetric Adjustment & Lower Amount (SAL): compared with Uncertain_OutcomeUnknown (i.e., 10 points), reciprocating more in Uncertain_Outcome8 (i.e., 16 points) and equivalently less in Uncertain_Outcome2 (i.e., 4 points), with the same amount of allocation in Uncertain_OutcomeUnknown but less total amount of allocation compared with Complete Adaptive Asymmetric Adjustment and Incomplete Adaptive Asymmetric Adjustment.
5. No Adjustment (NAD): reciprocating the same as Uncertain_OutcomeUnknown in both Uncertain_Outcome8 and in Uncertain_Outcome2 (i.e., 10 points in all conditions).
6. Selfish Asymmetric Adjustment (SAA): reciprocating significantly less in Uncertain_Outcome2 (i.e., 4 points) than in Uncertain_OutcomeUnknown (i.e., 10 points), and the same as Uncertain_OutcomeUnknown in Uncertain_Outcome8 (i.e., 10 points).

To be noted, with the exception of Symmetric Adjustment & Lower Amount, the amounts of the beneficiary’s allocation were fixed at 10 points in the Uncertain_OutcomeUnknown condition for all other patterns of reciprocity. Then for each pattern, the amounts of the beneficiary’s allocation in the Uncertain_Outcome2 condition and the Uncertain_Outcome8 condition were determined according to the type of the corresponding pattern and the number of points in the Uncertain_OutcomeUnknown condition. For Symmetric Adjustment & Higher Amount, the amount of the beneficiary’s allocation was set to 12 points in Uncertain_OutcomeUnknown condition, so that the total amount of allocation in Symmetric Adjustment & Higher Amount was comparable to Complete Adaptive Asymmetric Adjustment and Incomplete Adaptive Asymmetric Adjustment, which excluded the potential confounding caused by the different total amounts of allocation.

Specifically, in the questionnaire, each participant read the following background information: “In each round of an interpersonal game, the participant No. X was to receive 10 times of pain stimulation as a punishment for failing the task, and was randomly paired with a different anonymous co-player. The co-player (benefactor) decided whether to help No. X reduce the number of pain stimulation by 5 times, under uncertain cost of receiving a pain stimulation of either 2 times or 8 times, each with 50% probability. If the co-player decided to help, then the actual outcome of the cost that the co-player undertook, either 2 times or 8 times, was determined randomly by the computer and shown to both No. X and the co-player. The No. X had an opportunity to allocate some monetary points from own endowment (20 points) to the co-player as reciprocity. This decision might be made at one of the following three timepoints: (1) before the final outcome was ascertained (i.e., Uncertain_OutcomeUnknown condition);(2) after the final outcome was ascertained and the co-player actually undertook 2 times of pain stimulation (i.e., Uncertain_Outcome2 condition); (3) after the final outcome was ascertained and the co-player actually undertook 8 times of pain stimulation (i.e., Uncertain_Outcome8 condition).”

Then each participant would be presented with six scenarios describing the No. X’s amounts of allocations in the three conditions, corresponding to the above six allocation patterns, with the order of scenarios counterbalanced between participants. After reading each scenario, participants should rate the perceived morality (from -50, ‘very immoral,’ to 50, ‘very moral’), intention (from -50, ‘very unkind,’ to 50, ‘very kind’), stinginess (from 0, ‘very unstingy,’ to 100, ‘very stingy’), and utilitarianism (from 0, ‘very non-utilitarian,’ to 100, ‘very utilitarian’) of the person described. Additionally, they should rate their liking for this person (from -50, ‘very much disliked,’ to 50, ‘very liking’), and their willingness to be friends with this person (from 0, ‘no willingness,’ to 100, ‘very willing’). For example, for “Complete Adaptive Asymmetric Adjustment”, participants made ratings after seeing the following instructions: “One co-player decided to help No. X. If the allocation was made before the final outcome was ascertained, No. X decided to give the co-player 10 points. If the allocation was made after the final outcome was ascertained and the co-player actually undertook 2 times of pain stimulation, No. X decided to give the co-player 10 points. If the allocation was made after the final outcome was ascertained and the co-player actually undertook 8 times of pain stimulation, No. X decided to give the co-player 16 points. How would you make ratings about No .X?”

##### Model-free behavioral data analyses

Behavioral data analyses were all carried out in R 4.2.3 (https://www.r-project.org/). Bonferroni corrections were used for multiple comparison correction, and Greenhouse-Geisser corrections were used when the assumption of sphericity was violated.

##### Experiment 1

In Experiment 1, first, to reveal whether and how a beneficiary dynamically adjusted gratitude from facing an uncertain outcome (i.e., benefactor’s cost with a 50% chance of being either 2 or 8, expected to be 5 times of pain stimulation) to facing a final actual outcome of higher (i.e., 8) or lower (i.e., 2) benefactor’s cost than expected, we conducted one-way (Outcome in the Uncertain-to-Certain situation: Uncertain_Outcome2 vs. Uncertain_OutcomeUnknown vs. Uncertain_Outcome8) repeated-measure analyses of variance (ANOVA) for gratitude ratings. To further reveal how the direction of outcome change (from an uncertain cost with the expectation of 5 to final cost of 2 or to final cost of 8) influence the magnitude of dynamic adjustment, we subtracted the gratitude ratings in the Uncertain_Outcome2 and Uncertain_Outcome8 conditions from those in the Uncertain_OutcomeUnknown condition respectively, and took the absolute values as the indicators of the extent of the dynamic adjustment, and conducted Paired-samples *t*-tests.

Second, we aimed to examine whether and how the influence of actual outcomes of the benefactor’s cost (2 vs. 8) on the beneficiary’s gratitude responses differ between the Constantly-Certain situation and the Uncertain-to-Certain situation, despite the final outcomes of benefactor’s cost being equivalent in both situations, we combined the data of gratitude ratings in the Certain_Outcome2, Certain_Outcome8, Uncertain_Outcome2 and Uncertain_Outcome8 conditions, and performed 2 (Situation: Uncertain-to-Certain vs. Constantly-Certain) × 2 (Actual Outcome: 2 vs. 8) ANOVA for each variable respectively.

##### Experiment 2-4 and Experiment 7

The behavioral analyses of Experiment 2-4 and Experiment 7 were the same as Experiment 1, except that (1) In Experiments 4 and 7, we conducted analyses on participants’ monetary allocations instead of gratitude ratings; (2) in Experiment 3, we further compared the extent of “asymmetric adjustment” in gratitude-related responses between the context involving exogenous uncertainty in the benefactor’s cost and the context involving exogenous uncertainty in the beneficiary’s (participant’s) benefit by including context as an independent variable (see *SI Results*).

##### Experiment 5

With the participant’s monetary allocations as the dependent variable, we first conducted a 6 (Outcome magnitude: 2, 3, 4, 6, 7, vs. 8) × 2 (Situation: Uncertain-to-Certain vs. Constantly-Certain) repeated-measures ANOVA to examine whether and how the impact of the benefactor’s actual cost on the beneficiary’s allocations differed between the Constantly-Certain and Uncertain-to-Certain situations. Furthermore, a 3 (Uncertain-to-Certain condition: Uncertain 2-or-8 vs. 3-or-7 vs. 4-or-6) × 3 (Phase of monetary allocation: Uncertain_LowerOutcome vs. Uncertain_OutcomeUnknown vs. Uncertain_HigherOutcome) ANOVA was performed to investigate how various outcome combinations under uncertainty influenced the magnitude of dynamic adjustment in reciprocity from exogenous uncertainty to certainty. Finally, to examine whether there existed variations in the extent of the “asymmetric adjustment” across these Uncertain-to-Certain conditions with different outcome combinations, we calculated the magnitude of changes in reciprocity from the uncertain to the certain phase and submitted these change scores to an ANOVA, with the numeric manipulation of the Uncertain-to-Certain context (Uncertain 2-or-8 vs. 3-or-7 vs. 4-or-6) and the direction of change (from Uncertainty to Higher Outcome vs. from Uncertainty to Lower Outcome) as factors.

##### Experiment 6

Similar as Experiment 5, first, to examine whether and how the impact of the benefactor’s actual cost on the beneficiary’s allocations differed between the Constantly-Certain and Uncertain-to-Certain situations, and to investigate how various outcome combinations under uncertainty influenced the magnitude of dynamic adjustment in reciprocity from exogenous uncertainty to certainty, we conducted 6 (Situation: Uncertain-to-Certain 20%8 vs. Uncertain-to-Certain 40%8 vs. Uncertain-to-Certain 50%8 vs. Uncertain-to-Certain 60%8 vs. Uncertain-to-Certain 80%8 vs. Constantly-Certain) × 3 (Phase of monetary allocation: Uncertain_LowerOutcome vs. Uncertain_OutcomeUnknown/Uncertain_Outcome5 vs. Uncertain_HigherOutcome) ANOVA on monetary allocations. Second, to examine whether there existed variations in the extent of the “asymmetric adjustment” across these Uncertain-to-Certain conditions with different prior expectations, we calculated the magnitude of changes in reciprocity from the uncertain to the certain phase and submitted these change scores to an ANOVA, with the numeric manipulation of the Uncertain-to-Certain context (Uncertain 2-or-8 vs. 3-or-7 vs. 4-or-6) and the direction of change (from Uncertainty to Higher Outcome vs. from Uncertainty to Lower Outcome) as factors.

More importantly, we conducted computational modeling to formally distinguish between the “Differential PE Sensitivity” Hypothesis and the “Intrinsic Prosocial Bias” Hypothesis. For the values of changes in monetary allocations observed before and after the resolution of uncertainty, we fitted and compared 14 models that were built on the PE-based evaluation hypothesis (M1.1-M1.3), the “Intrinsic Prosocial Bias” Hypothesis (M2.1-M2.2), the combinations of these two mechanisms (M3.1-M3.6), as well as the baseline control models of outcome-based evaluation (M4.1-M4.2) and intention-based evaluation (M5.1), respectively. See Table S10 for details of all the models. See *SI Methods* for the details of model fitting and parameter recovery.

##### Experiment 8

To test whether the “asymmetric adjustment” in reciprocity during the exogenous uncertain-to-certain transition exhibited social adaptive function of increasing social acceptance and reputation, for each type of third-party ratings (perceived morality, intention, stinginess and utilitarianism, the liking for this person, and the willingness to be friends with this person), we performed one-way ANOVA on the rating values across the six scenarios of allocations patterns.

##### FMRI data acquisition and preprocessing of Experiment 1

Images were acquired through a 3T MRI scanner (Siemens Prisma; Siemens, Erlangen, Germany) with a 64-channel head coil. T2-weighted functional images were acquired using a single-shot T2*-weighted gradient-echo echo-planar imaging pulse sequence (TR = 1000 ms, TE = 32 ms, flip angle [FA] = 55°, each volume comprising 72 axial slices, matrix = 96 × 96, field of view [FoV] read = 192 mm, voxel size = 2 × 2 × 2 mm^3^). The fMRI data preprocessing and univariate analyses were conducted using the Statistical Parametric Mapping software SPM12 (Wellcome Trust Department of Cognitive Neurology, London, UK). Images were slice-time corrected (with the middle slice as the reference, i.e., the 36th slice), motion corrected, resampled to 2 mm × 2 mm × 2 mm isotropic voxels, and normalized to MNI space using the EPInorm approach in which functional images are aligned to an EPI template, which is then nonlinearly warped to stereotactic space^77^. Images were then spatially smoothed with a 5 mm FWHM Gaussian filter, and temporally filtered using a high-pass filter with a cutoff frequency of 1/128 Hz. T1-based normalization was not applied as the field maps necessary to adjust geometric distortion of EPI relative to the T1 images were not obtained^77^.

##### General linear model (GLM) analysis

A GLM analysis was conducted at the individual level (i.e., first-level analysis) in SPM12 to identify participants’ brain responses to the co-player’s decision of help or not help in each of the Uncertain_OutcomeUnknown, Uncertain_Outcome2, Uncertain_Outcome8, Certain_Outcome2, Certain_Outcome5, and Certain_Outcome8 conditions. In the GLM, we built a design matrix with separable run-specific partitions and with 12 key regressors (corresponding to the above 6 conditions in Help and NoHelp trials) in each run. All these 12 key regressors spanned from the presentation of the corresponding phase to the end of this event (3 s) (see *SI Methods*).

Regressors of no interest included Regrouping (onset of the Regrouping phase, 2-4 s), Situation information (onset of the Situation information phase, 2-4 s), Outcome_AfterAllocation (onset of the Outcome_Display phase of the trials in Uncertain_OutcomeUnknown condition, these brain responses were excluded from our main analysis due to the concern that the Allocation procedure might influence the subsequent processing of actual outcomes), Allocation (onset of the Allocation phase, spanning to the time that the participant finished allocation), and Miss_Allocation (onset of the Allocation phase of the trials in which participants failed to respond, 12 s). Six movement parameters were included as regressors of no interest. All regressors were convolved with a double gamma hemodynamic response function (HRF), and high-pass temporal filtering was applied with a default cutoff value of 128 s to eliminate low-frequency drifts. We defined 12 contrasts corresponding to the simple effects of the 12 key regressors.

##### Multivariate pattern analysis (MVPA)

MVPA was carried out in Python 3.8.10 using the NLTools package version 0.4.7 (https://nltools.org/)^78^. For each binary classification, we used the contrast images of the corresponding binary conditions from participants and linear Support Vector Machine (SVM)^54,55^ to train a whole-brain multivariate pattern classifier discriminating these two conditions (e.g., Uncertain_Outcome8 condition, coded as 1 vs. Uncertain_Outcome2 condition, coded as 0). SVM was conducted using linear kernel (regularization parameter C = 1), which has been suggested as a reasonable setting in multivariate pattern analysis^79^. With a leave-one-subject-out cross-validation (LOSO) method, we calculated the accuracy and significance of the SVM classifier using the forced-choice discrimination test^54,80,81^. Similar classification procedure was applied to all the following classification analyses. To be noted, first, during cross-validation, we ensured the independence of training and test data by holding out all images from the same participant together. Second, we compared the pattern expression values, calculated as the dot product of the vectorized activation image with the classifier weights, between two conditions within the same individual that was not part of the training sample. The condition with the higher pattern expression value was labeled as the 1 (e.g., Uncertain_Outcome8 condition), while the one with the lower value was labeled as 0 (e.g., Uncertain_Outcome2 condition)^54,80,81^.

Our behavioral and computational evidence consistently points to an asymmetric intrinsic prosocial bias, rather than a PE-based evaluation, as the primary driver of the “asymmetric adjustment” in reciprocity during the resolution of exogenous uncertainty. Under this mechanistic framework, the cognitive and neural representations triggered by the revelation of actual outcomes in the Uncertain-to-certain situation ought to diverge fundamentally from how identical outcomes are processed in the Constantly-Certain situation. From this perspective, we explored the unique neural representations in the Uncertain-to-Certain situation by comparing the neural representation of actual outcome in the Uncertain-to-Certain situation (Help_Uncertain_Outcome8 vs. Help_Uncertain_Outcome2 contrast maps) and that in the Constantly-Certain situation (Help_Certain_Outcome8 vs. Help_Certain_Outcome2 contrast maps).

###### Whole-brain multivariate classifications

We applied SVM to train a whole-brain multivariate pattern classifier discriminating Help_Uncertain_Outcome8 vs. Help_Uncertain_Outcome2 contrast maps (the Uncertain-to-Certain classifier), as well as a whole-brain multivariate pattern classifier discriminating Help_Certain_Outcome8 vs. Help_Certain_Outcome2 contrast maps (the Constantly-Certain classifier). Next, we conducted cross-situation classifications to examine whether there existed similar or different neural representations of the benefactor’s cost in these two situations at whole-brain level. To test whether the pattern for the Uncertain-to-Certain classifier could predict the two conditions in the Constantly-Certain situation, for each participant, we computing the dot-products (i.e., pattern expression values) of the contrast maps for Certain_Outcome8 and Certain_Outcome2 conditions based on the Uncertain-to-Certain classifier. Significantly higher pattern expression values in Certain_Outcome8 condition than in Certain_Outcome2 condition would indicate similar neural representations across situations, while insignificant difference would indicate differential neural representations. Similar analysis was conducted to test whether the Constantly-Certain classifier could predict the two conditions in the Uncertain-to-Certain situation.

###### Functional parcellation-based MVPA

We then aimed to search for specific brain regions that were specifically involved in the Constantly-Certain situation and in the Uncertain-to-Certain situation, respectively, within the 66 regions of interest (ROI) identified in the RSA. For each ROI, we applied SVM^54,55^ to train a multivariate pattern classifier discriminating Help_Uncertain_Outcome8 vs. Help_Uncertain_Outcome2 contrast maps, as well as a multivariate pattern classifier discriminating Help_Certain_Outcome8 vs. Help_Certain_Outcome2 contrast maps. Multi-tests were corrected using Bonferroni correction (i.e., *p* < 0.00076, two-tailed).

##### Neurosynth meta-analytical decoding

To investigate whether the processing of actual outcome in the Uncertain-to-Certain situation and the Constantly-Certain situation were associated with differential psychological components, we conducted meta-analytical decoding on the multivariate pattern maps for these two situations using the Neurosynth Image Decoder (neurosynth.org)^60^. This analysis allowed us to quantitatively evaluate the level of similarity between these two multivariate pattern maps (Fig. 4D) and each selected meta-analytical image generated by the Neurosynth database, indicated by the effect of spatial correlation (*r* value) between the two maps. If a term of psychological component was involved more in the Uncertain-to-Certain situation than in the Constantly-Certain situation, the similarity between this term and the multivariate pattern map for the Uncertain-to-Certain situation would be larger than that between this term and the multivariate pattern map for the Constantly-Certain situation, and vice versa. Referring to previous studies on social emotions in help-receiving situation^28,41^, Psychological terms were selected according to previous reviews on basic cognition (i.e., Imagine, Switching, Salience, Conflict, Memory, Attention, Cognitive control, Inhibition, Emotion, Anxiety, Fear, and Default mode)^82^, social cognition (Empathy, Theory of mind, Social, and Imitation)^83^ and decision-making (Reward, Punishment, Learning, Prediction error, Choice, and Outcome)^20^.

## Author Contributions

X.L., R.L., Y.H., X.Z., and X.G. designed the experiments; X.L., C.Y., and Y.N. implemented the study designs and collected the data; X.L., C.Y., R.L., Y.N., Y.F., and X.G. carried out the analyses; X.L., C.Y., R.L., Y.N., Y.F., Y.H., X.Z., and X.G. wrote the paper; and X.Z. and X.G. supervised the work. All authors provided critical revisions and approved the final paper for submission.

## Data and Code Availability

All data needed to evaluate the conclusions in the paper are present in the paper and/or the *Supplementary Materials*. Original materials will be made available in a trusted open-access repository (e.g., Github) after being accepted for publication.

## Acknowledgements

This work was supported by the National Natural Science Foundation of China (32571231, 32371094, X.G.), the Research Project of Shanghai Science and Technology Commission (20dz2260300, X.G. and X.Z.), and the Fundamental Research Funds for the Central Universities (X.G. and X.Z.).

## References

1. Fehr, E. & Gächter, S. Fairness and retaliation: The economics of reciprocity. Journal of Economic Perspectives 14, 159–182 (2000).

2. Henrich, J. & Muthukrishna, M. The origins and psychology of human cooperation. Annu. Rev. Psychol. 72, 207–240 (2021).

3. Malmendier, U., Te Velde, V. L. & Weber, R. A. Rethinking reciprocity. Annual Review of Economics 6, 849–874 (2014).

4. Nowak, M. A. Five rules for the evolution of cooperation. Science 314, 1560–1563 (2006).

5. Rand, D. G. & Nowak, M. A. Human cooperation. Trends in Cognitive Sciences 17, 413–425 (2013).

6. Schmid, L., Chatterjee, K., Hilbe, C. & Nowak, M. A. A unified framework of direct and indirect reciprocity. Nat Hum Behav 5, 1292–1302 (2021).

7. FeldmanHall, O. & Shenhav, A. Resolving uncertainty in a social world. Nat Hum Behav 3, 426–435 (2019).

8. Griffin, M. A. & Grote, G. When is more uncertainty better? A model of uncertainty regulation and effectiveness. AMR 45, 745–765 (2020).

9. Behrens, T. E. J., Hunt, L. T., Woolrich, M. W. & Rushworth, M. F. S. Associative learning of social value. Nature 456, 245–249 (2008).

10. Diaconescu, A. O. et al. Inferring on the intentions of others by hierarchical bayesian learning. PLoS Comput Biol 10, e1003810 (2014).

11. King-Casas, B. et al. Getting to know you: Reputation and trust in a two-person economic exchange. Science 308, 78–83 (2005).

12. Olsson, A., Knapska, E. & Lindström, B. The neural and computational systems of social learning. Nat Rev Neurosci 21, 197–212 (2020).

13. Ding, K. et al. The expectation-updating mechanism in gratitude: A predictive coding perspective. Emotion 25, 198–209 (2025).

14. Heffner, J., Son, J.-Y. & FeldmanHall, O. Emotion prediction errors guide socially adaptive behaviour. Nat Hum Behav 5, 1391–1401 (2021).

15. Vives, M.-L. & FeldmanHall, O. Tolerance to ambiguous uncertainty predicts prosocial behavior. Nat Commun 9, 2156 (2018).

16. Hilbe, C., Šimsa, Š., Chatterjee, K. & Nowak, M. A. Evolution of cooperation in stochastic games. Nature 559, 246–249 (2018).

17. Toyokawa, W., Whalen, A. & Laland, K. N. Social learning strategies regulate the wisdom and madness of interactive crowds. Nat Hum Behav 3, 183–193 (2019).

18. Downey, H. K. & Slocum, J. W. Uncertainty: Measures, research, and sources of variation. The Academy of Management Journal 18, 562–578 (1975).

19. Knight, F. Risk-Uncertainty-and-Profit. (Boston: Houghton Mifflin Company, 1921).

20. Ruff, C. C. & Fehr, E. The neurobiology of rewards and values in social decision making. Nat Rev Neurosci 15, 549–562 (2014).

21. Bavel, J. J. V. et al. Using social and behavioural science to support COVID-19 pandemic response. Nat Hum Behav 4, 460–471 (2020).

22. Fehr, E. On the economics and biology of trust. Journal of the European Economic Association 7, 235–266 (2009).

23. Gintis, H., Bowles, S., Boyd, R. & Fehr, E. Moral Sentiments and Material Interests: The Foundations of Cooperation in Economic Life. (The MIT Press, 2005).

24. Hallsworth, M. A manifesto for applying behavioural science. Nat Hum Behav 7, 310–322 (2023).

25. Kappes, A. et al. Uncertainty about the impact of social decisions increases prosocial behaviour. Nat Hum Behav 2, 573–580 (2018).

26. Chang, L. J., Smith, A., Dufwenberg, M. & Sanfey, A. G. Triangulating the neural, psychological, and economic bases of guilt aversion. Neuron 70, 560–572 (2011).

27. Fehr, E., Fischbacher, U. & Kosfeld, M. Neuroeconomic foundations of trust and social preferences: Initial evidence. American Economic Review 95, 346–351 (2005).

28. Gao, X. et al. The psychological, computational, and neural foundations of indebtedness. Nat Commun 15, 68 (2024).

29. Liao, R. et al. Efficiency recalibrates social-emotional trade-offs behind partner choice in direct reciprocity through intention-specific neural bases. Advanced Science 13, e16509 (2026).

30. Rilling, J. K. & Sanfey, A. G. The neuroscience of social decision-making. Annu. Rev. Psychol. 62, 23–48 (2011).

31. Sanfey, A. G. Social decision-making: Insights from game theory and neuroscience. Science 318, 598–602 (2007).

32. Van Baar, J. M., Chang, L. J. & Sanfey, A. G. The computational and neural substrates of moral strategies in social decision-making. Nat Commun 10, 1483 (2019).

33. Van Den Bos, W., Van Dijk, E., Westenberg, M., Rombouts, S. A. R. B. & Crone, E. A. What motivates repayment? Neural correlates of reciprocity in the trust game. Social Cognitive and Affective Neuroscience 4, 294–304 (2009).

34. Yu, H., Cai, Q., Shen, B., Gao, X. & Zhou, X. Neural substrates and social consequences of interpersonal gratitude: Intention matters. Emotion 17, 589–601 (2017).

35. Yu, H., Gao, X., Zhou, Y. & Zhou, X. Decomposing gratitude: Representation and integration of cognitive antecedents of gratitude in the brain. J. Neurosci. 38, 4886–4898 (2018).

36. Fehr, E. & Schmidt, K. M. Chapter 8 the economics of fairness, reciprocity and altruism – experimental evidence and new theories. in Foundations (eds Kolm, S.-C. & Ythier, J. M.) vol. 1 615–691 (Elsevier, 2006).

37. Falk, A. & Fischbacher, U. A theory of reciprocity. Games and Economic Behavior 54, 293–315 (2006).

38. McCabe, K. A., Rigdon, M. L. & Smith, V. L. Positive reciprocity and intentions in trust games. Journal of Economic Behavior & Organization 52, 267–275 (2003).

39. Nihonsugi, T., Ihara, A. & Haruno, M. Selective increase of intention-based economic decisions by noninvasive brain stimulation to the dorsolateral prefrontal cortex. J. Neurosci. 35, 3412–3419 (2015).

40. Tesser, A. Some determinants of gratitude. Journal of Personality and Social Psychology 9, 233–236 (1968).

41. Xiong, W. et al. Affective evaluation of others’ altruistic decisions under risk and ambiguity. NeuroImage 218, 116996 (2020).

42. Fox, G. R., Kaplan, J., Damasio, H. & Damasio, A. Neural correlates of gratitude. Front. Psychol. 6, (2015).

43. Kahneman, D. & Tversky, A. Prospect theory: An analysis of decision under risk. Econometrica 47, 263–291 (1979).

44. Ruggeri, K. et al. Replicating patterns of prospect theory for decision under risk. Nat Hum Behav 4, 622–633 (2020).

45. Tom, S. M., Fox, C. R., Trepel, C. & Poldrack, R. A. The neural basis of loss aversion in decision-making under risk. Science 315, 515–518 (2007).

46. Lefebvre, G., Lebreton, M., Meyniel, F., Bourgeois-Gironde, S. & Palminteri, S. Behavioural and neural characterization of optimistic reinforcement learning. Nat Hum Behav 1, 0067 (2017).

47. Mende-Siedlecki, P., Baron, S. G. & Todorov, A. Diagnostic value underlies asymmetric updating of impressions in the morality and ability domains. J. Neurosci. 33, 19406–19415 (2013).

48. Siegel, J. Z., Mathys, C., Rutledge, R. B. & Crockett, M. J. Beliefs about bad people are volatile. Nat Hum Behav 2, 750–756 (2018).

49. Haselton, M. G. & Nettle, D. The paranoid optimist: An integrative evolutionary model of cognitive biases. Pers Soc Psychol Rev 10, 47–66 (2006).

50. Marr, D. Vision: A Computational Investigation into the Human Representation and Processing of Visual Information. (MIT Press, 2010).

51. Yu, H., Gao, X., Shen, B., Hu, Y. & Zhou, X. A levels-of-analysis framework for studying social emotions. Nat Rev Psychol 3, 198–213 (2024).

52. Mørkbak, M. R., Olsen, S. B. & Campbell, D. Behavioral implications of providing real incentives in stated choice experiments. Journal of Economic Psychology 45, 102–116 (2014).

53. Trimmer, P. et al. Decision-making under uncertainty: Biases and bayesians. Animal cognition 14, 465–76 (2011).

54. Wager, T. D. et al. An fMRI-based neurologic signature of physical pain. New England Journal of Medicine 368, 1388–1397 (2013).

55. Hastie, T., Tibshirani, R. & Friedman, J. The Elements of Statistical Learning: Data Mining, Inference, and Prediction, Second Edition. (Springer, 2009).

56. Hu, Y., Gao, X., Yu, H., He, Z. & Zhou, X. Neuroscience of moral decision making. in Encyclopedia of Behavioral Neuroscience, 2nd edition 481–495 (Elsevier, 2022).

57. Koster-Hale, J. & Saxe, R. Theory of mind: A neural prediction problem. Neuron 79, 836–848 (2013).

58. Schaafsma, S. M., Pfaff, D. W., Spunt, R. P. & Adolphs, R. Deconstructing and reconstructing theory of mind. Trends in Cognitive Sciences 19, 65–72 (2015).

59. Schurz, M., Radua, J., Aichhorn, M., Richlan, F. & Perner, J. Fractionating theory of mind: A meta-analysis of functional brain imaging studies. Neuroscience & Biobehavioral Reviews 42, 9–34 (2014).

60. Yarkoni, T., Poldrack, R. A., Nichols, T. E., Van Essen, D. C. & Wager, T. D. Large-scale automated synthesis of human functional neuroimaging data. Nat Methods 8, 665–670 (2011).

61. Algoe, S. B., Fredrickson, B. L. & Gable, S. L. The social functions of the emotion of gratitude via expression. Emotion 13, 605–609 (2013).

62. McCullough, M. E., Kilpatrick, S. D., Emmons, R. A. & Larson, D. B. Is gratitude a moral affect? Psychological Bulletin 127, 249–266 (2001).

63. McCullough, M. E., Kimeldorf, M. B. & Cohen, A. D. An adaptation for altruism: The social causes, social effects, and social evolution of gratitude. Curr Dir Psychol Sci 17, 281–285 (2008).

64. Haselton, M. G. & Buss, D. M. Error management theory: A new perspective on biases in cross-sex mind reading. Journal of Personality and Social Psychology 78, 81–91 (2000).

65. Sears, D. O. The person-positivity bias. Journal of Personality and Social Psychology 44, 233–250 (1983).

66. Murray, S. L., Holmes, J. G. & Griffin, D. W. The benefits of positive illusions: Idealization and the construction of satisfaction in close relationships. Journal of Personality and Social Psychology 70, 79–98 (1996).

67. Van Overwalle, F. & Baetens, K. Understanding others’ actions and goals by mirror and mentalizing systems: A meta-analysis. NeuroImage 48, 564–584 (2009).

68. Baumgartner, T. Dorsolateral and ventromedial prefrontal cortex orchestrate normative choice.

69. González, B. & Chang, L. J. Computational models of mentalizing. in The Neural Basis of Mentalizing (eds Gilead, M. & Ochsner, K. N.) 299–315 (Springer International Publishing, Cham, 2021).

70. Hampton, A. N., Bossaerts, P. & O’Doherty, J. P. Neural correlates of mentalizing-related computations during strategic interactions in humans. Proc. Natl. Acad. Sci. U.S.A. 105, 6741–6746 (2008).

71. Le Petit, M. et al. Socially shared emotions shape the activity of the medial prefrontal cortex during inference of others’ emotional states. Cell Reports 45, 117506 (2026).

72. Wittmann, M. K. et al. Basis functions for complex social decisions in dorsomedial frontal cortex. Nature 641, 707–717 (2025).

73. Unoka, Z., Seres, I., Áspán, N. & Nikoletta, B. Trust game reveals restricted interpersonal transactions in patients with borderline personality disorder. Journal of personality disorders 23, 399–409 (2009).

74. Tsvetkova, M., Yasseri, T., Pescetelli, N. & Werner, T. A new sociology of humans and machines. Nat Hum Behav 8, 1864–1876 (2024).

75. Cheng, X. et al. The conceptual structure of human relationships across modern and historical cultures. Nat Hum Behav 9, 1162–1175 (2025).

76. Inui, K., Tran, T. D., Hoshiyama, M. & Kakigi, R. Preferential stimulation of adelta fibers by intra-epidermal needle electrode in humans. Pain 96, 247–252 (2002).

77. Calhoun, V. D. et al. The impact of T1 versus EPI spatial normalization templates for fMRI data analyses. Human Brain Mapping 38, 5331–5342 (2017).

78. Haynes, J.-D. & Rees, G. Decoding mental states from brain activity in humans. Nat Rev Neurosci 7, 523–534 (2006).

79. Varoquaux, G. et al. Assessing and tuning brain decoders: Cross-validation, caveats, and guidelines. NeuroImage 145, 166–179 (2017).

80. Chang, L. J., Gianaros, P. J., Manuck, S. B., Krishnan, A. & Wager, T. D. A sensitive and specific neural signature for picture-induced negative affect. PLoS Biol 13, e1002180 (2015).

81. Woo, C.-W. et al. Separate neural representations for physical pain and social rejection. Nat Commun 5, 5380 (2014).

82. Barrett, L. F. & Satpute, A. B. Large-scale brain networks in affective and social neuroscience: Towards an integrative functional architecture of the brain. Curr Opin Neurobiol 23, 361–372 (2013).

83. Adolphs, R. The social brain: Neural basis of social knowledge. Annu Rev Psychol 60, 693–716 (2009).

