## Supporting Information for "Computational Resilience in Human Reciprocity: An Asymmetric Intrinsic Prosocial Bias for Sustaining Cooperation under Exogenous Uncertainty"

**This PDF file includes:**

SI Methods
- Participants
- Model fitting and validations in Experiment 6
- Key regressors of fMRI general linear model (GLM) analysis in Experiment 7
- fMRI Representational similarity analysis (RSA) in Experiment 7

Supporting Tables

Supporting Figures

References for Supporting information

### SI Methods

**Participants**

For Experiment 1, 33 undergraduate and graduate Chinese Han students (23 females; 21.85 ± 1.64 years) were recruited from Shanghai, China, to complete a behavioral experiment at East China Normal University. All the participants were included in the data analysis.

For Experiment 2, 34 undergraduate and graduate Chinese Han students were recruited from Shanghai, China, to complete a behavioral experiment at East China Normal University. Three participants were excluded due to failing the comprehension test, leaving 31 participants (22 females; 21.4 ± 1.6 years) for data analysis.

For Experiment 3, 39 undergraduate and graduate Chinese Han students were recruited from Shanghai, China, to complete a behavioral experiment via a Chinese online experiment platform (NAODAO: www.naodao.com)[1]. Six participants were excluded due to failing the comprehension test, leaving 33 participants (16 females; 21.4 ± 2.0 years) for data analysis.

For Experiment 4, 37 undergraduate and graduate Chinese Han students (24 females; 20.62 ± 1.66 years) were recruited from Shanghai, China, to complete a behavioral experiment at East China Normal University. All the participants were included in the data analysis.

For Experiment 5, 43 undergraduate and graduate Chinese Han students were recruited from Shanghai, China, to participate in the behavioral experiment at East China Normal University. Five participants were excluded due to failing the comprehension test, leaving 38 participants (25 females; 20.45 ± 2.58 years) for data analysis.

For Experiment 6, 45 undergraduate and graduate Chinese Han students were recruited from Shanghai, China, to complete a behavioral experiment at East China Normal University. Two participants were excluded due to failing the comprehension test, leaving 43 participants (28 females; 20.95 ± 2.34 years) for data analysis.

For Experiment 7 (fMRI experiment), 49 undergraduate and graduate Chinese Han students were recruited from Shanghai, China, to participate in the fMRI experiment at East China Normal University. All participants were right-handed with normal or corrected-to-normal vision. Three participants were excluded from data analysis due to excessive head movements (>2mm of locomotion or >2° of rotation), leaving 46 participants (23 females; 21.7 ± 2.1 years) for data analysis.

For Experiment 8 (online questionnaire), 49 undergraduate and graduate Chinese Han students (21 females; 21.9 ± 2.0 years) were recruited from a Chinese online experiment platform (NAODAO: www.naodao.com) to complete an online questionnaire.

For all experiments, none of the participants reported any history of psychiatric, neurological, or cognitive disorders. All experiments were carried out in accordance with the Declaration of Helsinki and were approved by the Ethics Committee of East China Normal University. Informed written consent was obtained from each participant prior to participating.

#### Model fitting and validations in Experiment 6

*Hierarchical Bayesian model estimation*

We estimated the computational models using hierarchical Bayesian analysis (HBA)[2] implemented in the "rstan" package (version 2.32.7)[3] in R. Rstan uses Hamiltonian Monte Carlo sampling to approximate the full posterior distribution of model parameters. We used HBA rather than maximum-likelihood estimation because hierarchical models borrow information across participants and thereby regularize individual-level estimates. This shrinkage is particularly useful when individual behavioral data are noisy or sparse[4].

*Model Specification and Non-centered Parameterization*

The winning model used a piecewise structure to estimate behavioral sensitivity to the outcomes higher and lower than expected separately. Specifically, parameters were estimated for the PE > 0 and PE < 0 regions, allowing responses to positive and negative stimuli to vary independently.

To improve sampling efficiency and reduce the funnel geometry often observed in hierarchical models, we used a non-centered parameterization. For example, individual-level slopes $\alpha_i$ were derived as:

$$\alpha_i = \mu_\alpha + \sigma_\alpha \cdot z_{\alpha,i}, \; z_{\alpha,i} \sim \text{Normal}(0, 1)$$

with $\mu_\alpha$ and $\sigma_\alpha$ being the group-level mean and standard deviation, respectively and $z_{\alpha,i}$ denotes the individual-specific latent displacement.

*Robust Likelihood and Priors*

To reduce the influence of outlying behavioral observations, we used a Student's *t* likelihood. The likelihood included a degrees-of-freedom parameter, $\nu$, which allows heavier tails than a Gaussian likelihood.

We assigned weakly informative priors to group-level hyperparameters:
Means (μ): Normal (0, 50) for bias (intercepts) and Normal (0, 15) for slopes.

Scales (σ): Normal (0, 15) with a lower = 0 constraint, effectively forming a Half-Normal prior that regularizes the variance without imposing a non-zero mode.

*Sampling Efficiency and Convergence*

For each model, we ran four independent Markov chain Monte Carlo chains with 8,000 iterations per chain, including 4,000 warm-up iterations. This yielded 16,000 (= (8000 - 4000) × 4). Convergence was assessed using the Gelman-Rubin statistic ($\hat{R}$) and visual inspection of trace plots. For all parameters in the winning model, $\hat{R}$ values were ≤ 1.001, indicating excellent convergence (Table S13) [5].

*Model comparison and posterior predictive check (PPC)*

Model comparison. Candidate models were compared using the leave-one-out information criterion (LOOIC)[6]. LOOIC estimates expected out-of-sample predictive accuracy from the full posterior distribution and is therefore more appropriate for hierarchical Bayesian models than criteria based on point estimates (e.g., AIC or DIC). The model with the lowest LOOIC was selected as the winning model.

Posterior Predictive Check (PPC). We then evaluated the absolute fit of the winning model using PPCs[7] at three scales: Trial-wise, individual-wise, and grand-wise PPCs. At trial-wise and individual wise PPCs, we quantified the correspondence between observed behavior and synthetic data generated from the posterior predictive distribution. At the grand-wise PPC, we tested whether the observed mean behavior fell within the 95% highest-density interval (HDI) of the posterior predictive distribution. These checks assessed whether the model captured both individual-level variation and the overall behavioral pattern.

*Parameter recovery*

We assessed parameter identifiability by performing parameter recovery for the winning model.

First, we sampled group-level parameters from the prior-relevant parameter space of the winning model, such as a group-level mean ($\mu_{\alpha\_pos}$) and a group-level standard deviation ($\sigma_{\alpha\_pos}$) of the parameter $\alpha_{pos}$:

$$\alpha_{pos,i} \sim Normal(\mu_{\alpha\_pos}, \sigma_{\alpha\_pos})$$

The same procedure was applied to all group-level parameters in the winning model. We then generated synthetic behavioral datasets for 43 simulated participants performing the same task structure as in Experiment 6. For each simulated participant, dynamic adjustments in reciprocity ($\Delta \mathrm{Alloc}_i$) were sampled from the model likelihood conditional on that participant's latent parameters:

$$\Delta \mathrm{Alloc}_i \sim p(\Delta Alloc \mid \alpha_{pos}, \alpha_{neg}, \varepsilon_{pos}, \varepsilon_{neg})$$

Then, we refitted the winning model to the simulated data using the same HBA procedure applied to the empirical data. This yielded posterior estimates for both group-level parameters, such as $\mu'_{\alpha\_pos}$ and $\sigma'_{\alpha\_pos}$, and the individual level parameters, such as $\alpha'_{pos}$.

Finally, we compared whether the posterior distributions at both the group-level and the individual-level given the simulated data recovered the actual data generating parameters that were used to simulate those data.

**Key regressors of fMRI general linear model (GLM) analysis in Experiment 7**

- R1: Help_Uncertain_OutcomeUnknown, onset of the Outcome_Unknown phase of Help trials in the Uncertain_OutcomeUnknown condition, which captured the brain responses to the co-player's help under uncertainty in the Uncertain-to-Certain situation;
- R2: Help_Uncertain_Outcome2, onset of the Outcome_Display phase of Help trials in the Uncertain_Outcome2 condition, which captured the brain responses to the co-player's help after knowing the benefactor's actual cost was lower than expected in the Uncertain-to-Certain situation;
- R3: Help_Uncertain_Outcome8, onset of the Outcome_Display phase of Help trials in the Uncertain_Outcome8 condition, which captured the brain responses to the co-player's help after knowing the benefactor's actual cost was higher than expected in the Uncertain-to-Certain situation;
- R4: Help_Certain_Outcome2, onset of the Outcome_Display phase of Help trials in the Certain_Outcome2 condition, which captured the brain responses to the co-player's help under certain cost of 2 times in the Constantly-Certain situation;
- R5: Help_Certain_Outcome5, onset of the Outcome_Display phase of Help trials in the Certain_Outcome5 condition, which captured the brain responses to the co-player's help under certain cost of 5 times in the Constantly-Certain situation;
- R6: Help_Certain_Outcome8, onset of the Outcome_Display phase of Help trials in the Certain_Outcome8 condition, which captured the brain responses to the co-player's help under certain cost of 8 times in the Constantly-Certain situation.

The settings for R7 - R12, i.e., NoHelp_Uncertain_OutcomeUnknown, NoHelp_Uncertain_Outcome2, NoHelp_Uncertain_Outcome8, NoHelp_Certain_Outcome2, NoHelp_Certain_Outcome5, NoHelp_Certain_Outcome8, were the same as R1 - R6 respectively, but focused on the NoHelp trials.

**fMRI Representational similarity analysis (RSA) in Experiment 7**

The within-subject RSA was carried out in Python 3.8.10 using the NLTools package version 0.4.7 (https://nltools.org/) to search for the specific brain regions that were involved in different cognitive components of gratitude- and reciprocity-related processing, including the experience of gratitude indicated by gratitude ratings, the gratitude-induced reciprocity indicated by monetary allocation, the evaluation of kind intention indicated by perceived kind intention ratings, and the processing of benefactor's cost implemented in the experimental design, respectively. Each contrast image for each condition and each participant was divided into 200 parcels using a priori 200-parcel whole-brain parcellation based on meta-analytically functional co-activation of the Neurosynth database[8–10] (https://www.neurosynth.org/). The use of parcellation is less computationally demanding and exhibit higher homogeneity with functional neuroanatomy than the more conventional searchlight approach[8,10,11], and has been proven efficient in previous studies[8,10].

To search for brain regions involved in benefactor's cost related processing, for each parcel of each participant, we created a representational dissimilarity matrix (RDM) of neural activities using the pairwise correlation dissimilarity between each pair of the 12 key conditions (6 Help and 6 NoHelp conditions). Meanwhile, we constructed a behavioral RDM based on the values of benefactor's cost as implemented in the experimental design (Fig. S4A) by estimating the Euclidean distance between the corresponding cost values of each pair of the 12 key conditions (6 Help and 6 NoHelp conditions). Then, for each parcel and each participant, we applied Spearman's rank-order correlations on the lower triangle of the matrices to calculate the correlation between the parcel dissimilarity matrix and each behavioral dissimilarity matrix. For each parcel, we extracted the correlation coefficients ($\rho$ values) from all participants, conducted a Fisher $z$-transformation, and then conducted a one-sample sign permutation test to evaluate the association between the parcel dissimilarity matrix and each behavioral dissimilarity matrix at group-level. Multi-tests were corrected using Bonferroni correction (i.e., $p < 0.00025$, two-tailed). A similar procedure was adopted for searching for brain regions involved in intention-related and allocation-related processing, with the self-reported perceived kind intention and amount of monetary allocation, respectively, in the 12 key conditions (6 Help and 6 NoHelp conditions) as the inputs for the behavioral RDM (Fig. S4A). However, since participants made ratings on self-reported gratitude only regarding the six Help conditions, when searching for brain regions involved in gratitude-related processing, we constructed both the behavioral and neural RDMs using the data from these six conditions for each participant.

Moreover, previous studies[12–15] have consistently identified the association between gratitude and neural activities in ventral medial prefrontal cortex (vmPFC). Therefore, in addition to whole-brain parcellation based RSA, we conducted regions of interest (ROI) based analysis using the parcel ID 148 in the priori 200-parcel whole-brain parcellation of the Neurosynth database, which corresponding to the peak coordinate of vmPFC (MNI coordinate: 3, 44, 4) identified in Xiong et al. (2020)[13].

### Supporting Tables

**Table S1. Behavioral results in the Uncertain-to-Certain situation for Experiments 1-4 & 7.**

| Experiment | Independent Variable | Dependent Variable | Uncertain_ Outcome2 | Uncertain_ OutcomeUnknown | Uncertain_ Outcome8 | *df* | Main effect *F* | $\eta^2_{\text{partial}}$ |
|---|---|---|---|---|---|---|---|---|
| Experiment 1 | Benefactor's Cost | Gratitude | 75.00 (1.95) | 78.40 (1.72) | 85.05 (1.62) | (1.43, 45.69) | 43.83 *** | 0.58 |
| Experiment 2 | Beneficiary's Benefit | Gratitude | 72.51 (2.78) | 75.45 (2.23) | 86.42 (2.00) | (1.15, 34.47) | 27.21 *** | 0.48 |
| Experiment 3 | Benefactor's Cost Block | Gratitude | 70.89 (2.94) | 73.58 (2.83) | 87.61 (1.71) | (1.57, 50.22) | 31.78 *** | 0.20 |
| | Beneficiary's Benefit Block | Gratitude | 73.14 (2.45) | 75.25 (2.46) | 85.26 (2.13) | (1.71, 54.76) | 23.52 *** | 0.42 |
| Experiment 4 | Benefactor's Cost | Allocation | 7.74 (0.68) | 9.08 (0.70) | 11.13 (0.72) | (1.36, 48.90) | 69.37 *** | 0.66 |
| Experiment 7 | Benefactor's Cost | Allocation | 8.84 (0.52) | 9.43 (0.49) | 10.95 (0.55) | (1.51, 67.78) | 61.82 *** | 0.58 |

Note: (1) Values in Uncertain_Outcome2, Uncertain_OutcomeUnknown, and Uncertain Outocme8 conditions are presented as Mean (SE). (2) * $p < 0.05$, ** $p < 0.01$, and *** $p < 0.001$.

**Table S2. The effects of Uncertain-to-Certain Transition on beneficiaries' gratitude ratings and monetary allocations in the Uncertain-to-Certain situation for Experiments 1-4 & 7.**

| Experiment | Independent Variable | Dependent Variable | Lower than Expectation | Higher than Expectation | *df* | Paired-samples *t* | Cohen's *d* |
|---|---|---|---|---|---|---|---|
| Experiment 1 | Benefactor's Cost | Gratitude | 3.74 (0.69) | 6.84 (1.04) | 32 | 2.65 * | 0.46 |
| Experiment 2 | Beneficiary's Benefit | Gratitude | 5.52 (0.89) | 11.62 (1.62) | 30 | 3.93 *** | 0.71 |
| Experiment 3 | Benefactor's Cost Block | Gratitude | 5.68 (1.30) | 14.10 (2.43) | 32 | 3.29 ** | 0.57 |
| | Beneficiary's Benefit Block | Gratitude | 6.85 (1.11) | 10.74 (1.64) | | 2.60 * | 0.45 |
| Experiment 4 | Benefactor's Cost | Allocation | 1.39 (0.22) | 2.07 (0.24) | 36 | 2.43 * | 0.40 |
| Experiment 7 | Benefactor's Cost | Allocation | 0.80 (0.14) | 1.52 (0.17) | 45 | 3.68 *** | 0.54 |

Note: (1) The value of "Lower than Expectation" = |Uncertain_Outcome2 – Uncertain_OutcomeUnknown|, The value of "Higher than Expectation" = |Uncertain_Outcome8 – Uncertain_OutcomeUnknown|. Values of "Lower than Expectation" and "Higher than Expectation" are presented as Mean (SE). (2) * $p < 0.05$, ** $p < 0.01$, and *** $p < 0.001$.

**Table S3. The gratitude ratings in response to the actual outcomes of benefactor's cost in the Constantly-Certain situation and in the Uncertain-to-Certain situation for Experiment 1 and 7**

| Dependent Variable | Situation | Outcome8 | Outcome2 | *df* | Paired-samples *t* | Cohen's *d* |
|---|---|---|---|---|---|---|
| Gratitude | Uncertain-to-Certain | 85.05 (2.75) | 75.00 (1.95) | 32 | 7.36 *** | 0.81 |
| | Constantly-Certain | 94.60 (1.13) | 59.75 (3.19) | 32 | 11.41 *** | 2.80 |
| Allocation | Uncertain-to-Certain | 8.84 (0.52) | 10.95 (0.55) | 45 | 8.60 *** | 1.01 |
| | Constantly-Certain | 7.02 (0.49) | 12.72 (0.59) | 45 | 13.09 *** | 2.72 |

Note: (1) Values of gratitude ratings in Uncertain_Outcome8, Uncertain_Outcome2, Certain_Outcome8, and Certain_Outcome2 conditions are presented as Mean (SE). (2) * $p < 0.05$, ** $p < 0.01$, and *** $p < 0.001$.

**Table S4. The descriptive statistics and simple effects of monetary allocation in the Numeric Uncertain-to-Certain conditions of Experiment 5**

| Situation | LowerOutcome | OutcomeUnknown | HigherOutcome | *df* | Main effect *F* | $\eta^2_{\text{partial}}$ |
|---|---|---|---|---|---|---|
| Uncertain 4-or-6 | 41.08 (2.28) | 42.90 (2.32) | 47.12 (2.59) | | 21.17 *** | 0.53 |
| Uncertain 3-or-7 | 40.39 (2.71) | 45.82 (2.76) | 53.32 (2.89) | (2, 37) | 23.13 *** | 0.56 |
| Uncertain 2-or-8 | 40.53 (3.31) | 48.74 (3.20) | 59.99 (3.54) | | 25.49 *** | 0.58 |

Note: (1) Values of monetary allocation in LowerOutcome, OutcomeUnknown, and HigherOutcome conditions are presented as Mean (SE). (2) * $p < 0.05$, ** $p < 0.01$, and *** $p < 0.001$.

**Table S5. The extent of adjustment of monetary allocation in different Numeric Uncertain-to-Certain conditions in Experiment 5**

| Situation | From Uncertainty to Lower Outcome | From Uncertainty to Lower Outcome |
|---|---|---|
| Uncertain46 | 2.34 (0.41) | 4.45 (0.65) |
| Uncertain37 | 5.54 (1.05) | 7.91 (1.16) |
| Uncertain28 | 8.18 (1.47) | 11.68 (1.59) |

Note: (1) The value of monetary allocation in “From Uncertainty to Lower Outcome” = |LowerOutcome –OutcomeUnknown|, The value of “From Uncertainty to Higher Outcome” = |HigherOutcome –OutcomeUnknown|. Values of From Uncertainty to Lower Outcome and From Uncertainty to Lower Outcome conditions are presented as Mean (SE).

**Table S6. The descriptive statistics of monetary allocation in Constantly-Certain and Uncertain-to-Certain conditions in Experiment 5**

| | Uncertain-to-Certain | Constantly-Certain | *df* | *t* | Cohen's *d* |
|---|---|---|---|---|---|
| Outcome2 | 40.53 (3.31) | 26.61 (2.49) | 37 | 5.93 *** | 1.18 |
| Outcome3 | 40.39 (2.71) | 32.21 (2.20) | | 5.40 *** | 0.69 |
| Outcome4 | 41.08 (2.28) | 37.91 (2.17) | | 3.94 *** | 0.27 |
| Outcome6 | 47.12 (2.59) | 51.52 (2.77) | | -4.82 *** | -0.38 |
| Outcome7 | 53.32 (2.89) | 59.89 (3.23) | | -5.39 *** | -0.56 |
| Outcome8 | 59.99 (3.54) | 66.86 (3.31) | | -4.53 *** | -0.58 |

Note: (1) Values of monetary allocation in all conditions are presented as Mean (SE). (2) * $p < 0.05$, ** $p < 0.01$, and *** $p < 0.001$.

**Table S7. The participants' expectations regarding the final actual outcome under different experimental conditions in Experiments 2-7**

| Situation | Mean (SE) | *df* | *t* | *p* | Cohen's *d* |
|---|---|---|---|---|---|
| Experiment 2 | | | | | |
| Uncertain28 (Benefit) | 4.97 (0.22) | 30 | -0.14 | 0.887 | -0.03 |
| Experiment 3 | | | | | |
| Uncertain28 (Cost) | 5.15 (0.09) | 32 | 1.72 | 0.096 | 0.30 |
| Uncertain28 (Benefit) | 5.33 (0.21) | 32 | 1.61 | 0.117 | 0.28 |
| Experiment 4 | | | | | |
| Uncertain28 (Cost) | 5.08 (0.11) | 36 | 0.77 | 0.446 | 0.13 |
| Experiment 5 | | | | | |
| Uncertain28 (Cost) | 5.05 (0.21) | 37 | 0.25 | 0.806 | 0.04 |
| Uncertain37 (Cost) | 5.03 (0.18) | 37 | 0.14 | 0.886 | 0.02 |
| Uncertain46 (Cost) | 5.11 (0.18) | 37 | 0.60 | 0.554 | 0.10 |
| Experiment 6 | | | | | |
| 100%2 (Cost) | 2.02 (0.22) | 42 | 0.10 (vs. 2.0) | 0.918 | 0.02 |
| 100%5 (Cost) | 5.17 (1.41) | 42 | 0.77 (vs. 5.0) | 0.444 | 0.12 |
| 100%8 (Cost) | 7.61 (2.44) | 42 | -1.02 (vs. 8.0) | 0.312 | -0.16 |
| 20%8 (Cost) | 3.23 (0.20) | 42 | 0.16 (vs. 3.2) | 0.873 | 0.02 |
| 40%8 (Cost) | 4.26 (0.15) | 42 | -0.97 (vs. 4.4) | 0.339 | -0.15 |
| 50%8 (Cost) | 5.26 (0.15) | 42 | 1.67 (vs. 5.0) | 0.102 | 0.26 |
| 60%8 (Cost) | 5.88 (0.20) | 42 | 1.42 (vs. 5.6) | 0.164 | 0.22 |
| 80%8 (Cost) | 6.77 (0.25) | 42 | -0.13 (vs. 6.8) | 0.897 | -0.02 |
| Experiment 7 | | | | | |
| Uncertain28 (Cost) | 5.09 (0.20) | 45 | 0.44 | 0.664 | 0.06 |

**Table S8. The descriptive statistics and simple effects of monetary allocation in the probabilistic Uncertain-to-Certain experimental conditions of Experiment 6**

| Situation | Outcome2 | OutcomeUnknown / Outcome5 | Outcome8 | *df* | Main effect *F* | $\eta^2_{\text{partial}}$ |
|---|---|---|---|---|---|---|
| Uncertain-to-Certain 20%8 | 31.62 (2.68) | 32.90 (2.74) | 43.56 (2.95) | | 30.67 *** | 0.59 |
| Uncertain-to-Certain 40%8 | 32.90 (2.74) | 39.05 (2.62) | 43.56 (2.95) | | 26.28 *** | 0.56 |
| Uncertain-to-Certain 50%8 | 40.07 (2.72) | 44.34 (2.62) | 53.07 (2.61) | 42 | 35.34 *** | 0.63 |
| Uncertain-to-Certain 60%8 | 43.71 (2.92) | 48.27 (2.80) | 55.66 (2.73) | | 33.66 *** | 0.62 |
| Uncertain-to-Certain 80%8 | 52.94 (3.43) | 57.95 (3.16) | 64.85 (3.10) | | 30.21 *** | 0.59 |
| Constantly-Certain | 29.51 (2.77) | 47.91 (2.34) | 74.16 (3.11) | | 119.45 *** | 0.85 |

Note: (1) Values of monetary allocation in all conditions are presented as Mean (SE). (2) * $p < 0.05$, ** $p < 0.01$, and *** $p < 0.001$.

**Table S9. The extent of dynamic adjustment of monetary allocation in different PE values for each condition in Experiment 6**

| \|PE\| | Lower than Expectation | Higher than Expectation |
|---|---|---|
| \|PE\| = 1.2 | 2.82 (0.48) | 6.98 (1.04) |
| \|PE\| = 2.4 | 3.90 (0.65) | 7.39 (1.14) |
| \|PE\| = 3.0 | 5.04 (0.80) | 8.85 (1.23) |
| \|PE\| = 3.6 | 5.34 (0.87) | 8.62 (1.27) |
| \|PE\| = 4.8 | 6.05 (1.02) | 10.86 (1.50) |

Note: The value of "Lower than Expectation" = |Uncertain_LowerOutcome – Uncertain_OutcomeUnknown|, The value of "Higher than Expectation" = |Uncertain_HigherOutcome – Uncertain_OutcomeUnknown|. Values of "Lower than Expectation" and "Higher than Expectation" are presented as Mean (SE).

**Table S10. Computational models and model comparison results of Experiment 6**

| Model | Description | Formula | *nPars* | *LOOIC* | *ΔLOOIC* | *Weight* |
|---|---|---|---|---|---|---|
| PE-based evaluation | | | | | | |
| M1.1 | Consistent PE slope | $Allocation_{post} = Allocation_{pre} + \alpha \cdot PE$ | 1 | 17637.81 | 899.02 | 0 |
| M1.2 | Asymmetric PE slopes | $Allocation_{post} = \begin{cases} Allocation_{pre} + \alpha_{pos} \cdot PE & if\ PE > 0 \\ Allocation_{pre} + \alpha_{neg} \cdot PE & if\ PE < 0 \end{cases}$ | 2 | 16999.91 | 261.12 | 0 |
| M1.3 | Power-law scaling of \|PE\| | $Allocation_{post} = \begin{cases} Allocation_{pre} + \alpha \cdot \lvert PE \rvert^{\gamma} & if\ PE > 0 \\ Allocation_{pre} - \alpha \cdot \lvert PE \rvert^{\gamma} & if\ PE < 0 \end{cases}$ | 2 | 17589.80 | 851.01 | 0 |
| Intrinsic bias model | | | | | | |
| M2.1 | Only global bias | $Allocation_{post} = Allocation_{pre} + Bias$ | 1 | 18768.29 | 2029.51 | 0 |
| M2.2 | Only asymmetric bias | $Allocation_{post} = \begin{cases} Allocation_{pre} + Bias_{pos} & if\ PE > 0 \\ Allocation_{pre} + Bias_{neg} & if\ PE < 0 \end{cases}$ | 2 | 17167.22 | 428.43 | 0 |
| PE-based evaluation with intrinsic bias | | | | | | |
| M3.1 | Consistent PE slope with global bias | $Allocation_{post} = Allocation_{pre} + \alpha \cdot PE + Bias$ | 2 | 17019.23 | 280.44 | 0.004 |
| M3.2 | Consistent PE slope with PE-sign-dependent bias | $Allocation_{post} = \begin{cases} Allocation_{pre} + \alpha \cdot PE + Bias & if\ PE > 0 \\ Allocation_{pre} + \alpha \cdot PE - Bias & if\ PE < 0 \end{cases}$ | 2 | 17575.4 | 836.61 | 0 |
| M3.3 | Asymmetric PE slopes with global bias | $Allocation_{post} = \begin{cases} Allocation_{pre} + \alpha_{pos} \cdot PE + Bias & if\ PE > 0 \\ Allocation_{pre} + \alpha_{neg} \cdot PE + Bias & if\ PE < 0 \end{cases}$ | 3 | 16952.02 | 213.23 | 0 |
| M3.4 | Asymmetric PE slopes with PE-sign-dependent bias | $Allocation_{post} = \begin{cases} Allocation_{pre} + \alpha_{pos} \cdot PE + Bias & if\ PE > 0 \\ Allocation_{pre} + \alpha_{neg} \cdot PE - Bias & if\ PE < 0 \end{cases}$ | 3 | 16856.45 | 117.66 | 0 |
| M3.5 | Consistent PE slope with asymmetric bias | $Allocation_{post} = \begin{cases} Allocation_{pre} + \alpha \cdot PE + Bias_{pos} & if\ PE > 0 \\ Allocation_{pre} + \alpha \cdot PE + Bias_{neg} & if\ PE < 0 \end{cases}$ | 3 | 16834.64 | 95.85 | 0.001 |
| M3.6 | Asymmetric PE slopes with asymmetric bias | $Allocation_{post} = \begin{cases} Allocation_{pre} + \alpha_{pos} \cdot PE + Bias_{pos} & if\ PE > 0 \\ Allocation_{pre} + \alpha_{neg} \cdot PE + Bias_{neg} & if\ PE < 0 \end{cases}$ | 4 | 16738.79 | 0.00 | 0.999 |
| Outcome-based evaluation | | | | | | |
| M4.1 | Consistent Outcome slope | $Allocation_{post} = Allocation_{pre} + \beta \cdot Outcome$ | 1 | 19155.90 | 2417.11 | 0 |
| M4.2 | Power-law scaling of Outcome | $Allocation_{post} = Allocation_{pre} + \beta \cdot Outcome^{\gamma}$ | 2 | 17163.41 | 424.62 | 0 |
| Intention-based evaluation | | | | | | |
| M5.1 | Depending on the expected benefactor's cost (i.e., the perceived cost the benefactor willing to undertake) | $Allocation_{post} = Allocation_{pre}$ | 0 | 18756.63 | 2017.84 | 0 |

**Table S11. Description statistics (mean and 95% HDI) and parameter recovery results of the parameters of the winning model M3.6**

| Parameters | Description | Mean and 95% HDI | Parameter Recovery | | | |
|---|---|---|---|---|---|---|
| | | | Pearson's *r* | *df* | *t* | 95% CI |
| $\alpha_{pos}$ | Positive slope | 3.37 [2.40, 4.30] | 0.98 | 42 | 8.44 *** | [0.65, 0.89] |
| $Bias_{pos}$ | Positive bias | 4.70 [3.70, 5.62] | 0.98 | | 33.81 *** | [0.97, 0.99] |
| $\alpha_{neg}$ | Negative slope | 3.08 [2.27, 3.89] | 0.77 | | 7.69 *** | [0.61, 0.87] |
| $Bias_{neg}$ | Negative bias | -0.74 [-1.51, 0.00] | 0.80 | | 28.35 *** | [0.95,0.99] |

Note: *** $p < 0.001$.

**Table S12. Description statistics (mean and 95% HDI) and parameter recovery results of the parameters of the winning model M3.6**

| Parameters | Positive (PE > 0) | Negative (PE < 0) | *df* | *t* | Cohen's *d* |
|---|---|---|---|---|---|
| $\alpha$ | 3.37 (0.93) | 3.08 (0.88) | 42 | 0.27 n.s. | 0.04 |
| $Bias$ | 4.70 (0.96) | -0.74 (0.31) | | 5.34 *** | 0.81 |
| $\lvert Bias\rvert$ | 5.63 (0.83) | 1.68 (0.21) | | 5.05 *** | 0.77 |

Note: (1) Values of parameters in all conditions are presented as Mean (SE). *** $p < 0.001$, n.s. $p > 0.05$.

**Table S13. $\hat{R}$ values of all parameters in the winning model M3.6**

| | $\mu_{\beta\_pos}$ | $\sigma_{\beta\_pos}$ | $\mu_{\beta\_neg}$ | $\sigma_{\beta\_neg}$ | $\mu_{\varepsilon\_pos}$ | $\sigma_{\varepsilon\_pos}$ | $\mu_{\varepsilon\_neg}$ | $\sigma_{\varepsilon\_neg}$ |
|---|---|---|---|---|---|---|---|---|
| $\hat{R}$ | 1.000 | 1.000 | 1.000 | 1.001 | 1.000 | 1.000 | 1.000 | 1.000 |

Note: $\hat{R}$ values of all parameters in the winning model M3.6 were close to 1.0 (smaller than 1.1 at most in the current study), which indicated adequate convergence.

**Table S14. The third-party evaluation of different beneficiary-reciprocity patterns in the Uncertain-to-Certain situation**

| Mean (SE) | **CAAA** | **IAAA** | SAM | SAL | NAD | SAA | (df1, df2) | Main effect *F* |
|---|---|---|---|---|---|---|---|---|
| Morality | **29.22(1.73) [a]** | **27.43(2.05) [a]** | 20.14(2.39) [b] | 11.12(3.26) [c] | 12.46(2.84) [bc] | -5.98(3.36) [d] | (3.29, 158.03) | 33.34*** |
| Willingness to be friend | **71.35(2.72) [a]** | **67.04(2.64) [a]** | 64.12(3.21) [a] | 51.14(3.10) [b] | 50.18(3.17) [b] | 31.16(3.58) [c] | (3.46, 166.30) | 28.32*** |
| Intention | **29.86(1.97) [a]** | **26.82(2.01) [a]** | 19.29(2.63) [b] | 10.92(2.77) [c] | 11.20(2.53) [bc] | -5.47(3.14) [d] | (3.51, 168.26) | 39.24*** |
| Impression | **30.06(1.81) [a]** | **26.43(2.09) [a]** | 17.35(2.44) [b] | 7.22(3.06) [c] | 12.47(2.63) [bc] | -5.55(3.34) [d] | (3.04, 145.72) | 35.19*** |
| Stinginess | **18.22(2.62) [a]** | **23.06(2.75) [a]** | 26.57(2.95) [a] | 40.49(3.76) [b] | 40.94(3.53) [b] | 61.63(3.73) [c] | (3.68, 176.51) | 34.22*** |
| Utilitarianism | **29.65(3.57) [a]** | **30.98(2.74) [a]** | 38.08(3.25) [a] | 49.86(3.72) [b] | 34.45(3.42) [a] | 61.06(3.46) [c] | (3.78, 181.39) | 20.46*** |

Note: CAAA = Complete Adaptive Asymmetric Adjustment; IAAA = Incomplete Adaptive Asymmetric Adjustment; SAM = Symmetric Adjustment & More Amount; SAL = Symmetric Adjustment & Less Amount; NAD = No Adjustment; SAA = Selfish Asymmetric Adjustment. Different letters above the statistical values on the right represent Compact Letter Display (CLD) groupings, indicating significant differences among groups in post hoc comparisons (corrected $p < 0.05$).

* $p < 0.05$, ** $p < 0.01$, and *** $p < 0.001$.

**Table S15. Trials setting in Experiment 1-4 & 7**

| | | Uncertain-to-Certain | | | Constantly-Certain | | | Total |
|---|---|---|---|---|---|---|---|---|
| | | Uncertain_ Outcome2 | Uncertain_ OutcomeUnknown | Uncertain_ Outcome8 | Certain_ Outcome2 | Certain_ Outcome5 | Certain_ Outcome8 | |
| Experiment 1 | Cost-Gratitude | 6 Help + 3 NoHelp | 12 Help + 6 NoHelp | 6 Help + 3 NoHelp | 12 Help + 6 NoHelp | 12 Help + 6 NoHelp | 12 Help + 6 NoHelp | 90 |
| Experiment 2 | Benefit-Gratitude | 6 Help + 3 NoHelp | 12 Help + 6 NoHelp | 6 Help + 3 NoHelp | 6 Help + 3 NoHelp | 6 Help + 3 NoHelp | 6 Help + 3 NoHelp | 63 |
| Experiment 3 | Cost-Gratitude | 6 Help + 3 NoHelp | 12 Help + 6 NoHelp | 6 Help + 3 NoHelp | 6 Help + 3 NoHelp | 6 Help + 3 NoHelp | 6 Help + 3 NoHelp | 126 |
| | Benefit-Gratitude | 6 Help + 3 NoHelp | 12 Help + 6 NoHelp | 6 Help + 3 NoHelp | 6 Help + 3 NoHelp | 6 Help + 3 NoHelp | 6 Help + 3 NoHelp | |
| Experiment 4 | Cost-Allocation | 8 Help + 4 NoHelp | 16 Help + 8 NoHelp | 8 Help + 4 NoHelp | 8 Help + 4 NoHelp | 8 Help + 4 NoHelp | 8 Help + 4 NoHelp | 84 |
| Experiment 7 | Cost-Allocation | 12 Help + 6 NoHelp | 24 Help + 6 NoHelp | 12 Help + 6 NoHelp | 12 Help + 6 NoHelp | 12 Help + 6 NoHelp | 12 Help + 6 NoHelp | 126 |

**Table S16. Trials setting in Experiment 5 & 6**

| ID | Experiment 5 | | Experiment 6 | |
|---|---|---|---|---|
| 1 | Uncertain46_ LowerOutcome | 6 Help + 3 NoHelp | 100%8 | 6 Help + 3 NoHelp |
| 2 | Uncertain46_ OutcomeUnknown | 6 Help + 3 NoHelp | 100%5 | 6 Help + 3 NoHelp |
| 3 | Uncertain46_ HigherOutcome | 6 Help + 3 NoHelp | 100%2 | 6 Help + 3 NoHelp |
| 4 | Uncertain46_ LowerOutcome | 6 Help + 3 NoHelp | 80%8_ Outcome2 | 6 Help + 3 NoHelp |
| 5 | Uncertain46_ OutcomeUnknown | 6 Help + 3 NoHelp | 80%8_ OutcomeUnknown | 6 Help + 3 NoHelp |
| 6 | Uncertain46_ HigherOutcome | 6 Help + 3 NoHelp | 80%8_ Outcome8 | 6 Help + 3 NoHelp |
| 7 | Uncertain46_ LowerOutcome | 6 Help + 3 NoHelp | 60%8_ Outcome2 | 6 Help + 3 NoHelp |
| 8 | Uncertain46_ OutcomeUnknown | 6 Help + 3 NoHelp | 60%8_ OutcomeUnknown | 6 Help + 3 NoHelp |
| 9 | Uncertain46_ HigherOutcome | 6 Help + 3 NoHelp | 60%8_ Outcome8 | 6 Help + 3 NoHelp |
| 10 | Certain_Outcome2 | 6 Help + 3 NoHelp | 50%8_ Outcome2 | 6 Help + 3 NoHelp |
| 11 | Certain_Outcome3 | 6 Help + 3 NoHelp | 50%8_ OutcomeUnknown | 6 Help + 3 NoHelp |
| 12 | Certain_Outcome4 | 6 Help + 3 NoHelp | 50%8_ Outcome8 | 6 Help + 3 NoHelp |
| 13 | Certain_Outcome5 | 6 Help + 3 NoHelp | 40%8_ Outcome2 | 6 Help + 3 NoHelp |
| 14 | Certain_Outcome6 | 6 Help + 3 NoHelp | 40%8_ OutcomeUnknown | 6 Help + 3 NoHelp |
| 15 | Certain_Outcome7 | 6 Help + 3 NoHelp | 40%8_ Outcome8 | 6 Help + 3 NoHelp |
| 16 | Certain_Outcome8 | 6 Help + 3 NoHelp | 20%8_ Outcome2 | 6 Help + 3 NoHelp |
| 17 | | | 20%8_ OutcomeUnknown | 6 Help + 3 NoHelp |
| 18 | | | 20%8_ Outcome8 | 6 Help + 3 NoHelp |
| Total | | 144 | | 162 |

### Supporting Figures

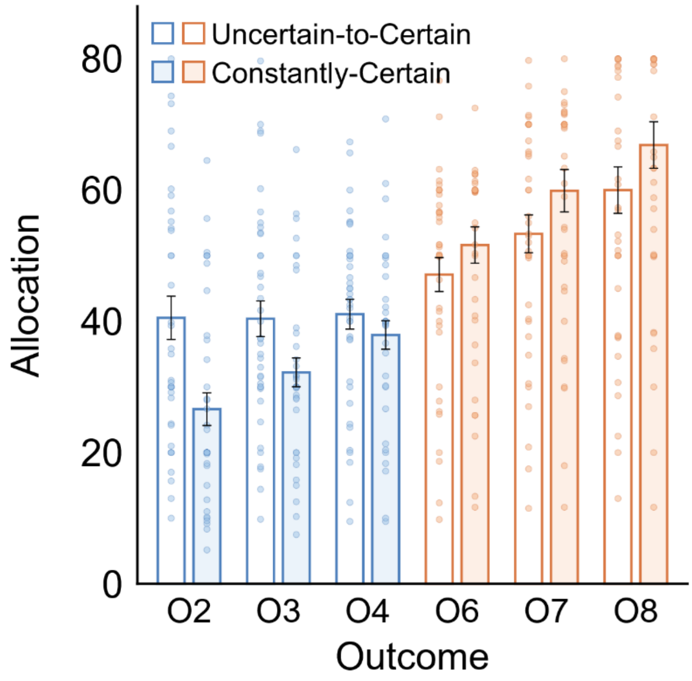


**Fig. S1 | Comparison of monetary allocations between the Uncertain-to-Certain and Constantly-Certain situations in Experiment 5.** Participants' monetary allocations across six different outcome magnitudes (2, 3, 4, 6, 7, and 8) are compared between the two situations (n = 38). A 6 (Outcome magnitude) × 2 (Situation) repeated-measures ANOVA revealed a significant interaction effect, indicating that the main effect of outcome magnitude on reciprocity was significantly attenuated in the Uncertain-to-Certain situation compared to the Constantly-Certain situation. Error bars represent standard errors.

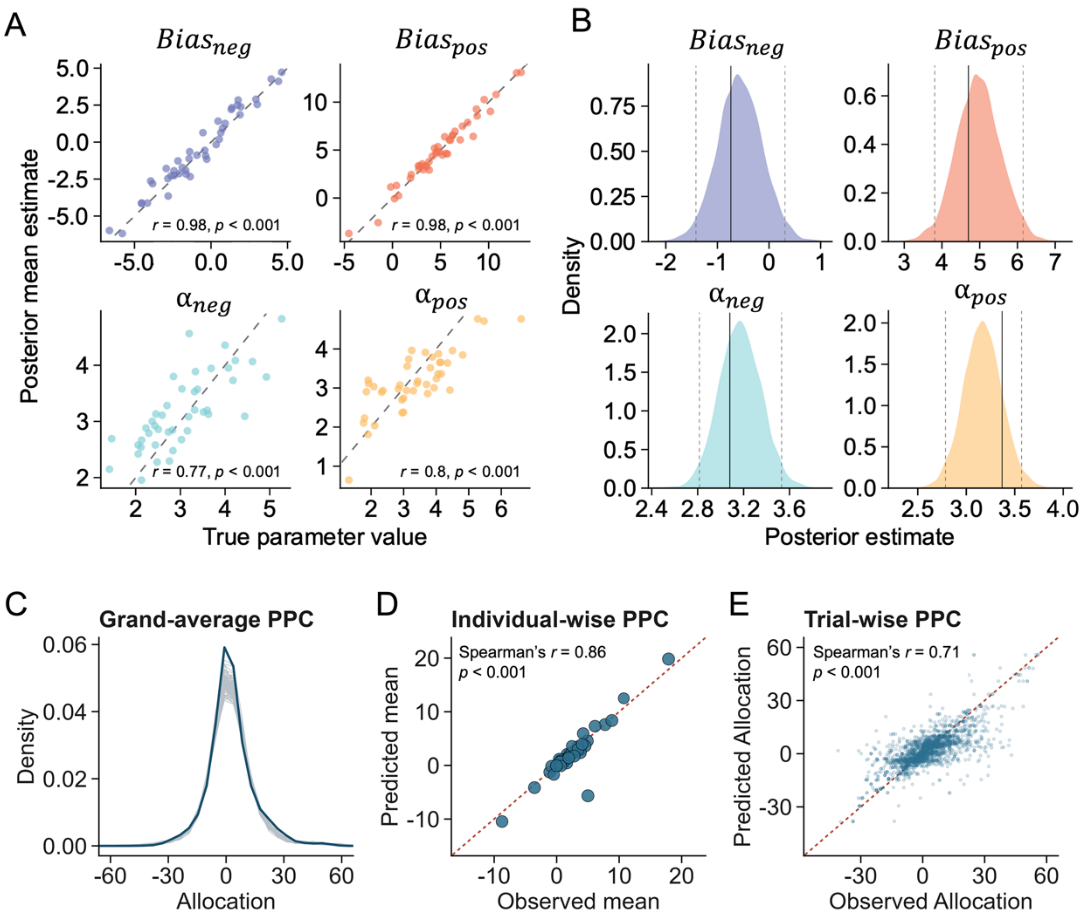


**Fig. S2 | Parameter recovery of the winning mode and posterior predictive check of the winning model in Experiment 6. (A-B)** All parameters could be accurately and selectively recovered, showing proper identifiability of model parameters: (C) for the individual level, true parameters (x-axis) and estimated parameters (y-axis) were well correlated. (B) for the group-level, true parameters (the black lines) falling within 95% HDI of each parameter's posterior density (n = 43). **(C-E)** Posterior predictive check of the winning model. (C) Grand-wise posterior predicative check. The density plot displays the distribution of observed adjustment-extent of reciprocity (y, represented by the thick dark line) against 100 sets of simulated adjustment-extent of reciprocity ($y_{rep}$, represented by the thin blue lines) generated from the model's posterior distributions. The high degree of overlap between the observed and predicted distributions indicates that the model successfully captures the central tendency and the overall variance of the empirical data at the aggregate level. (D) Individual-wise posterior predicative check. The actual individual data well correlated by the predicted data. (E) Trial-wise posterior predicative check. The actual trial-by-trial data well correlated by the predicted data.

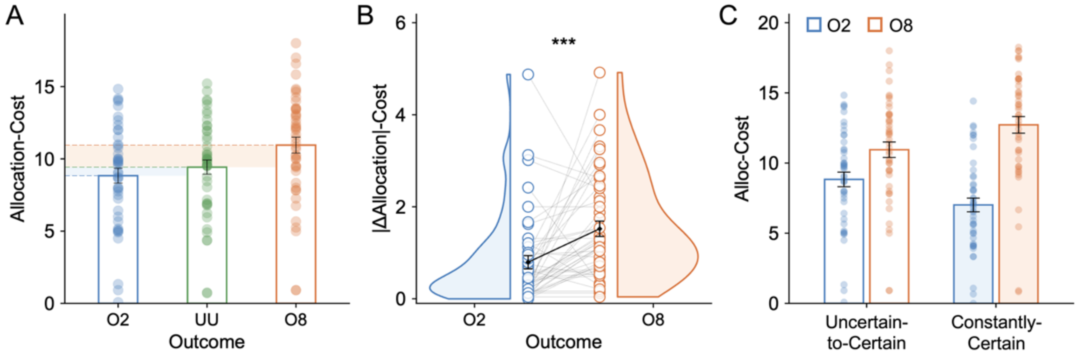


**Fig. S3 | The "asymmetric adjustment" in reciprocity under exogenous uncertainty in Experiment 7. (A)** Participants' monetary allocations for different actual benefactor costs (O2: Outcome = 2; UU: Uncertain_OutcomeUnknown; O8: Outcome = 8) in the Uncertain-to-Certain situation during the fMRI experiment (n = 46). **(B)** The absolute magnitude of adjustments in monetary allocations (|△Allocation|) when transitioning from uncertainty to an actual outcome of 2 or 8. Consistent with the findings from behavioral experiments, participants robustly exhibited the "asymmetric adjustment" in their reciprocity, where the adjustment was significantly larger when the outcome exceeded expectations than when it fell short. **(C)** Comparison of monetary allocations between the Uncertain-to-Certain and Constantly-Certain situations, demonstrating differential sensitivity to outcomes between the two situations during the transition from uncertainty to certainty. ***$p < 0.001$. Error bars represent standard errors.

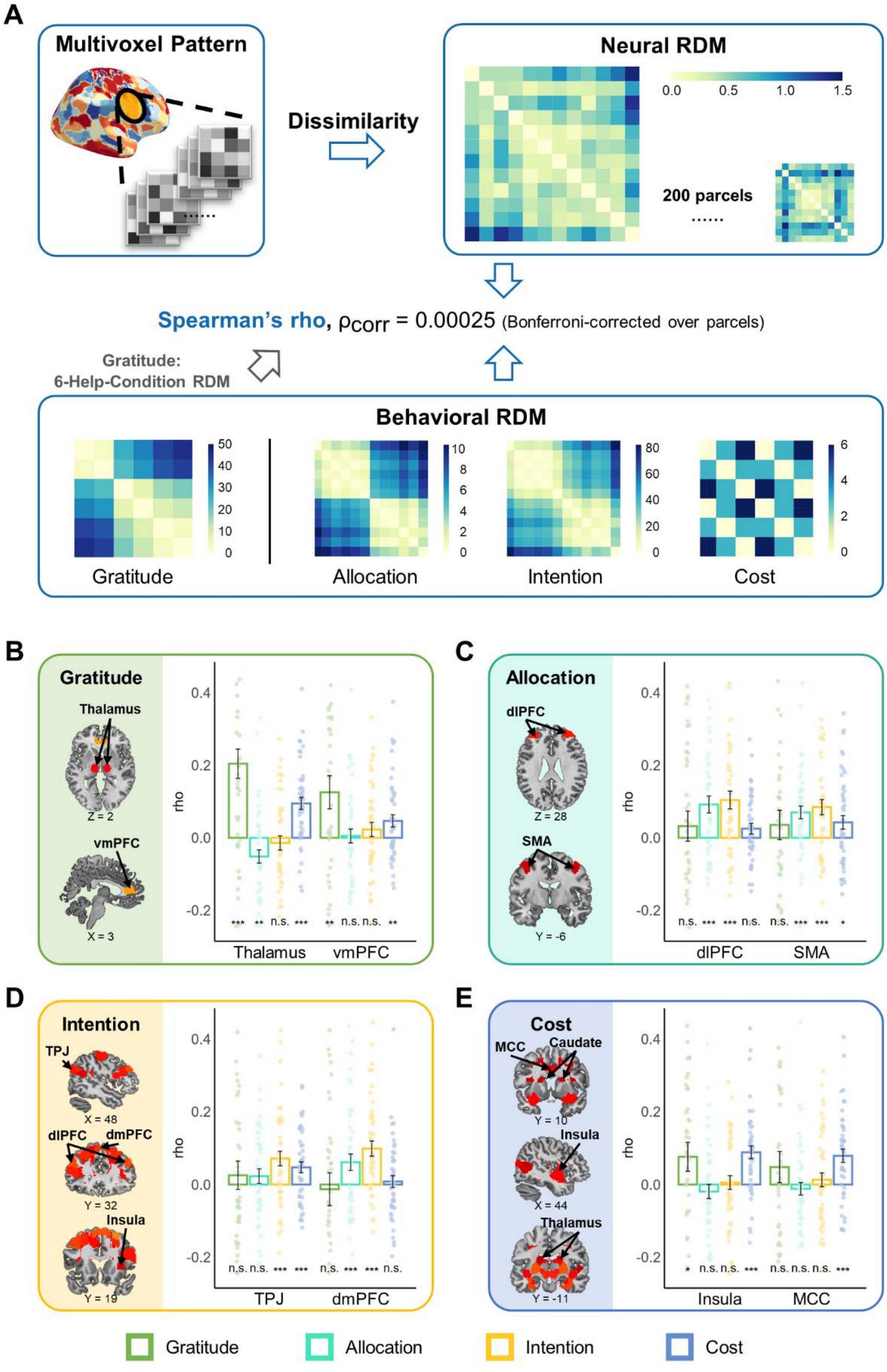


**Fig. S4 | Within-subject RSA of fMRI data in Experiment 7. (A)** At the neural level, each contrast image for each condition and each participant were divided into

200 parcels using a priori 200-parcel whole-brain parcellation[8–10]. For each parcel of each participant, we created a RDM of neural activities using the pairwise correlation dissimilarity between each pair of conditions. At behavioral level, we constructed the behavioral representational dissimilarity matrixes (RDM) for each of the behavioral indicators of these four cognitive components (gratitude rating, monetary allocation, kind intention rating, and benefactor's cost) respectively by estimating the Euclidean distance between the corresponding values of each pair of conditions. Then, for each parcel and each participant, we estimated the correlation between the neural RDM and each behavioral RDM using the Spearman rank-ordered correlation. For each parcel, we extracted the correlation coefficients ($\rho$ values) from all participants, conducted a Fisher $z$-transformation, and then conducted a one-sample sign permutation test to evaluate the association between the parcel dissimilarity matrix and each behavioral dissimilarity matrix at group-level. Multi-tests were corrected using Bonferroni correction (i.e., $p < 0.00025$, two-tailed). **(B-E)** Results showed that, for self-reported gratitude, we only identified one brain parcel located in the thalamus after whole-brain correction (B). Given that previous studies have identified that crucial role of vmPFC in gratitude processing[12–18,18–20], we further conducted ROI-based analysis using the parcel corresponding to the peak coordinate of vmPFC (MNI coordinate: 3, 44, 4) identified in Xiong et al. (2020)[13]. As expected, we discovered that the multivoxel activity pattern of this brain parcel was significantly correlated with the gratitude RDM ($r = 0.15$, $p = 0.007$). For monetary allocation, we identified two significant brain parcels, located in dorsolateral prefrontal cortex (dlPFC) and supplementary motor area (SMA) (C). For perceived kind intention ratings, we found forty significant brain parcels, including dmPFC, dlPFC, right insula, and bilateral temporo-parietal junction (TPJ) (D). For benefactor's cost, we observed twenty-six significant brain parcels, which were mainly located in the subcortical nucleus, including bilateral insula, bilateral striatum, hippocampus, midcingulate cortex (MCC), and thalamus (E). Error bars represent the SEs; significance: * $p < 0.05$; ** $p < 0.01$; *** $p < 0.001$.

#### References


1. Cheng, X. *et al.* The conceptual structure of human relationships across modern and historical cultures. *Nat Hum Behav* **9**, 1162–1175 (2025).
2. Gelman, A., Carlin, J. B., Stern, H. S. & Rubin, D. B. *Bayesian Data Analysis*. (Chapman and Hall/CRC, New York, 1995).
3. Carpenter, B. *et al.* Stan: A Probabilistic Programming Language. *J Stat Softw* **76**, 1 (2017).
4. Ahn, W.-Y., Haines, N. & Zhang, L. Revealing neurocomputational mechanisms of reinforcement learning and decision-making with the hBayesDM package. *Comput Psychiatr* **1**, 24–57 (2017).
5. Gelman, A. & Rubin, D. B. Inference from Iterative Simulation Using Multiple Sequences. *Statistical Science* **7**, 457–472 (1992).
6. Vehtari, A., Gelman, A. & Gabry, J. Practical Bayesian model evaluation using leave-one-out cross-validation and WAIC. *Stat Comput* **27**, 1413–1432 (2017).
7. Zhang, L., Lengersdorff, L., Mikus, N., Gläscher, J. & Lamm, C. Using reinforcement learning models in social neuroscience: Frameworks, pitfalls and suggestions of best practices. *Social Cognitive and Affective Neuroscience* **15**, 695–707 (2020).
8. Chang, L. J. *et al.* Endogenous variation in ventromedial prefrontal cortex state dynamics during naturalistic viewing reflects affective experience. *Sci. Adv.* **7**, eabf7129 (2021).
9. de la Vega, A., Chang, L. J., Banich, M. T., Wager, T. D. & Yarkoni, T. Large-scale meta-analysis of human medial frontal cortex reveals tripartite functional organization. *J Neurosci* **36**, 6553–6562 (2016).
10. Van Baar, J. M., Chang, L. J. & Sanfey, A. G. The computational and neural substrates of moral strategies in social decision-making. *Nat Commun* **10**, 1483 (2019).
11. Craddock, R. C., James, G. A., Holtzheimer, P. E., Hu, X. P. & Mayberg, H. S. A whole brain fMRI atlas generated via spatially constrained spectral clustering. *Hum Brain Mapp* **33**, 1914–1928 (2012).
12. Fox, G. R., Kaplan, J., Damasio, H. & Damasio, A. Neural correlates of gratitude. *Front. Psychol.* **6**, (2015).
13. Xiong, W. *et al.* Affective evaluation of others' altruistic decisions under risk and ambiguity. *NeuroImage* **218**, 116996 (2020).
14. Yu, H., Cai, Q., Shen, B., Gao, X. & Zhou, X. Neural substrates and social consequences of interpersonal gratitude: Intention matters. *Emotion* **17**, 589–601 (2017).
15. Yu, H., Gao, X., Zhou, Y. & Zhou, X. Decomposing gratitude: Representation and integration of cognitive antecedents of gratitude in the brain. *J. Neurosci.* **38**, 4886–4898 (2018).
16. Gao, X. *et al.* The psychological, computational, and neural foundations of indebtedness. *Nat Commun* **15**, 68 (2024).

17. Karns, C. M., Moore, W. E. & Mayr, U. The cultivation of pure altruism via gratitude: A functional MRI study of change with gratitude practice. *Front. Hum. Neurosci.* **11**, 599 (2017).

18. Kini, P., Wong, J., McInnis, S., Gabana, N. & Brown, J. W. The effects of gratitude expression on neural activity. *NeuroImage* **128**, 1–10 (2016).

19. Liu, G. *et al.* Neural responses to intention and benefit appraisal are critical in distinguishing gratitude and joy. *Sci Rep* **10**, 7864 (2020).

20. Zahn, R. *et al.* The neural basis of human social values: Evidence from functional MRI. *Cerebral Cortex* **19**, 276–283 (2009).
